# MINT Block: multivariate integration across independent studies to identify multi-omics signatures

**DOI:** 10.64898/2026.10.13.758954

**Authors:** David P. Hirst, Marko Terzin, Sean M. Burnard, Kim-Anh Lê Cao

## Abstract

Integrative multi-omics analyses can identify molecular signatures spanning multiple layers of a biological system. However, signatures derived from datasets with small sample sizes may not generalise. Combining data from multiple independent studies can increase the available sample size, but may also introduce unwanted variation arising from study-specific differences. Existing approaches typically address this variation sequentially by correcting the data before downstream model fitting, which can inadvertently distort the biological signals of interest. We introduce MINT Block, a framework that integrates multi-omics data across independent studies while simultaneously accounting for study-specific unwanted variation. MINT Block supports multivariate regression and classification analyses, enabling the identification of generalisable multi-omics signatures that capture relationships between different types of molecules and predict sample phenotypes. We evaluated MINT Block using bulk and single-cell multi-omics datasets from studies of human disease and marine biology. Across these applications, we showed that MINT Block identified stable, interpretable molecular signatures and improves prediction accuracy compared with existing sequential approaches.

## 1 Background

In recent years, the fields of biology and medicine have benefited from the development of high-throughput technologies that generate omics data (genomics, transcriptomics, etc.) by profiling molecules of the same type across a set of biological samples [1, 2]. Omics data are analysed with a variety of aims, including biomarker discovery [3], inferring relationships between molecules [4], and fitting classification and survival models [5]. Two important considerations in these analyses are that a) each type of omics data represents one of multiple layers in a biological system, and b) omics datasets typically contain a limited number of observations relative to the number of variables. These considerations can be addressed by the integrative analysis of multiple omics datasets. In our previous work, we have referred to these forms of integrative analysis as N-integration, P-integration, and NP-integration [6]. We use N-integration to refer to the integration of different types of omics data, generated for the same *N* samples or individuals. We use P-integration to refer to the integration of observations from independent studies for the same *P* omics variables, while NP-integration combines N- and P-integration.

Different types of omics represent different interacting layers of a biological system, so N-integration can provide more comprehensive insights than independent single-omics analyses [7, 8, 9]. N-integration can be challenging, however, as omics datasets differ in dimensionality, data type, and the sources and levels of variability. To address these challenges, N-integration strategies often rely on dimensionality reduction [9, 10].

We previously introduced an N-integration framework, based on Projection to Latent Structures (PLS) [11]. PLS is a dimensionality reduction approach that builds linear transformations to infer latent components, resulting in highly interpretable outputs [10, 12]. Our N-integration framework, which includes DIABLO [13], encompasses methods for multivariate regression and classification analyses that infer components with maximal covariance across different omics datasets. Covariance based N-integration methods have been shown to perform well in several benchmarking studies [14, 15, 9]

P-integration is motivated by the cost and difficulty of generating omics data, which often result in small sample sizes and limit the ability to draw meaningful, generalisable conclusions [16, 17, 18]. Researchers can increase the number of observations available for analysis by combining data from multiple studies. However, combining studies can introduce unwanted variation arising from differences in experimental design, sample collection, laboratory protocols and other extraneous factors [19, 5]. Unwanted variation can compromise reproducibility and generalisability, making it important to account for this variation in statistical analyses [20, 21]. Several methods have been developed for this purpose, many originally to address *batch effects* arising when samples are processed in different experimental batches within a study.

For many omics data analysis tasks, the only available option for mitigating unwanted variation is a sequential approach, in which variation is first removed and the *corrected* data are then used as input to the analysis [22, 23]. A common approach, as employed by ComBat [24], is to include known extraneous factors as covariates in a linear model and remove their estimated effects. Other methods, such as RUV-III [25], use lower-dimensional representations, while general normalisation methods may also alleviate unwanted variation associated with study or batch membership [23, 26]. Most methods for removing unwanted variation have been designed for bulk transcriptomics data, although exceptions include Harmony for single-cell RNA-seq data [27], and PLSDA-batch for microbiome data [28].

A limitation of sequential approaches is that correcting the data may either inflate biological signal or inadvertently remove it, thereby affecting false positive rates or statistical power. Thus, when possible, a simultaneous approach is often recommended, in which unwanted variation is not explicitly removed from the data but is instead accounted for during model fitting [19, 20, 29]. Methods for detecting differential expression, such as DESeq2 [30] and limma [31] are examples of simultaneous approaches. Another example is our P-Integration framework MINT [22], which encompasses methods for multivariate regression and classification analyses that account for study membership, and has been used in diverse research applications including human health [32, 33], biofuels [34, 35], and marine biology [36, 37]. A limitation of MINT is that it is only applicable to the integration of at most two omics types; similarly, other methods for addressing unwanted variation were designed for single-omics data. As such, there is a need for NP-integration methods that simultaneously account for unwanted variation.

To address the limitations of current N- and P-integration approaches for omics data, we present MINT Block, a novel NP-integration framework. MINT Block enables the integrative analysis of multiple types of omics data and/or other biological data, from multiple independent studies, for regression or classification analyses. We first present an overview of the framework and implementation, and then benchmark MINT Block against established sequential approaches, using multi-study, multi-omics and microbiome datasets from three different biological contexts. Finally, we present biological insights obtained from the application of MINT Block in these contexts.

## 2 Results

### 2.1 Overview of the MINT Block NP-integration framework

We first outline the data structure, analytical settings and outputs of the MINT Block framework, before evaluating its performance across three case studies and examining the biological signatures identified in each application (Figure 1).

**Figure 1.**
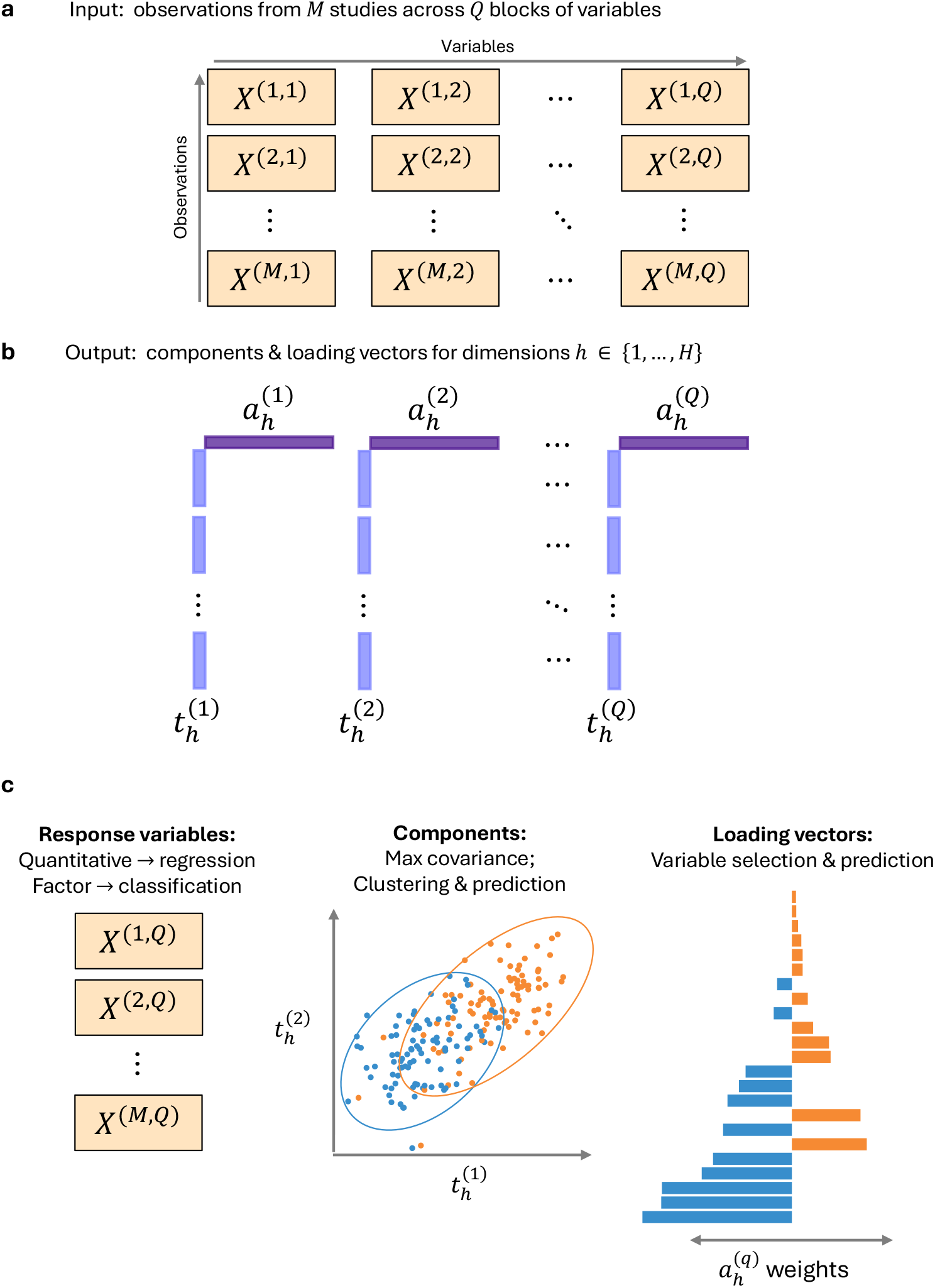
MINT Block jointly integrates multiple data blocks across independent studies while accounting for study-specific variation. **a)** The input comprises observations from *M* independent studies measured across *Q* blocks of variables. **b)** For each dimension *h* = 1, …, *H* and block *q*, MINT Block estimates a sparse loading vector 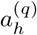 and corresponding component 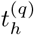. Before estimating the next dimension, the algorithm deflates each data matrix to obtain a residual matrix. Together, the *H* components provide a lower-dimensional representation of the dataset. **c)** MINT Block performs multivariate regression for quantitative response(s), or classification for a categorical response, by maximising co-variance among the block-specific components. The components support visualisation, clustering and prediction, and the loading vectors support variable selection and prediction.

We use the following notation. Consider a dataset *X* formed by combining observations from *M* ≥ 2 independent studies, each contributing the same *Q* ≥ 3 omics data blocks (at least two explanatory blocks, and one response block). In this paper, we refer to each data type as a *block*. We use *X*^(m,q)^ to denote the (*N*_m_ × *P*_q_) data matrix for study *m* and block *q*, where *N*_m_ is the number of observations in study *m* and *P*_q_ is the number of variables in block *q* and may range from one to thousands.

We present MINT Block for supervised analysis of *X* in either a regression or classification setting, where one of the blocks contains one or more response variables and the remaining blocks contain explanatory variables. For quantitative responses, MINT Block performs multivariate regression analysis (e.g. GBR dataset, see Section 2.2). For a categorical response indicating class membership, MINT Block performs classification (discriminant) analysis (e.g. the AML and T2D datasets, see Section 2.2).

MINT Block computes latent components and loading vectors for each of *H* dimensions, maximising the covariance among components across the different blocks while accounting for study-specific unwanted variation (Figure 1). The components are lower-dimensional representations of the input observations that can be used for visualisation, clustering and other downstream tasks. The loading vectors enable the selection of variables that covary across the blocks, providing insights into relationships within and between omics and identifying variables that discriminate between classes. Additionally, the components and loading vectors can be used to compute prediction coefficients for new data. For a detailed description of the MINT Block framework, see Section 4.1

### 2.2 Case studies

We evaluated the performance of MINT Block in multivariate regression and classification analyses, using datasets from three case studies, each comprising at least three blocks of omics or other biological data from multiple independent studies.

#### AML case study

We evaluated MINT Block classification using a single-cell multi-omics dataset comprising 184 acute myeloid leukemia (AML) cells profiled using scNMT-seq [38], a method that profiles DNA methylation, chromatin accessibility and gene expression (Figure 2a). For the DNA methylation and chromatin accessibility data, we restricted the analysis to sites located within CpG islands (CGIs). Each cell was treated with either azacytidine (AZA) or decitabine (DAC), which are DNA hypomethylating agents used in the treatment of AML. The resulting dataset consisted of four data blocks: DNA methylation at CGI sites (2,153 variables), chromatin accessibility at CGI sites (1,793 variables), RNA expression counts (4,975 variables), and a categorical response indicating the treatment received by each cell. The AML data were generated in three batches, which we treated as three independent studies in our analysis. See Section 4.2 for details on data acquisition and pre-processing, and Supplementary Table S1 for sample counts by treatment and batch.

**Figure 2.**
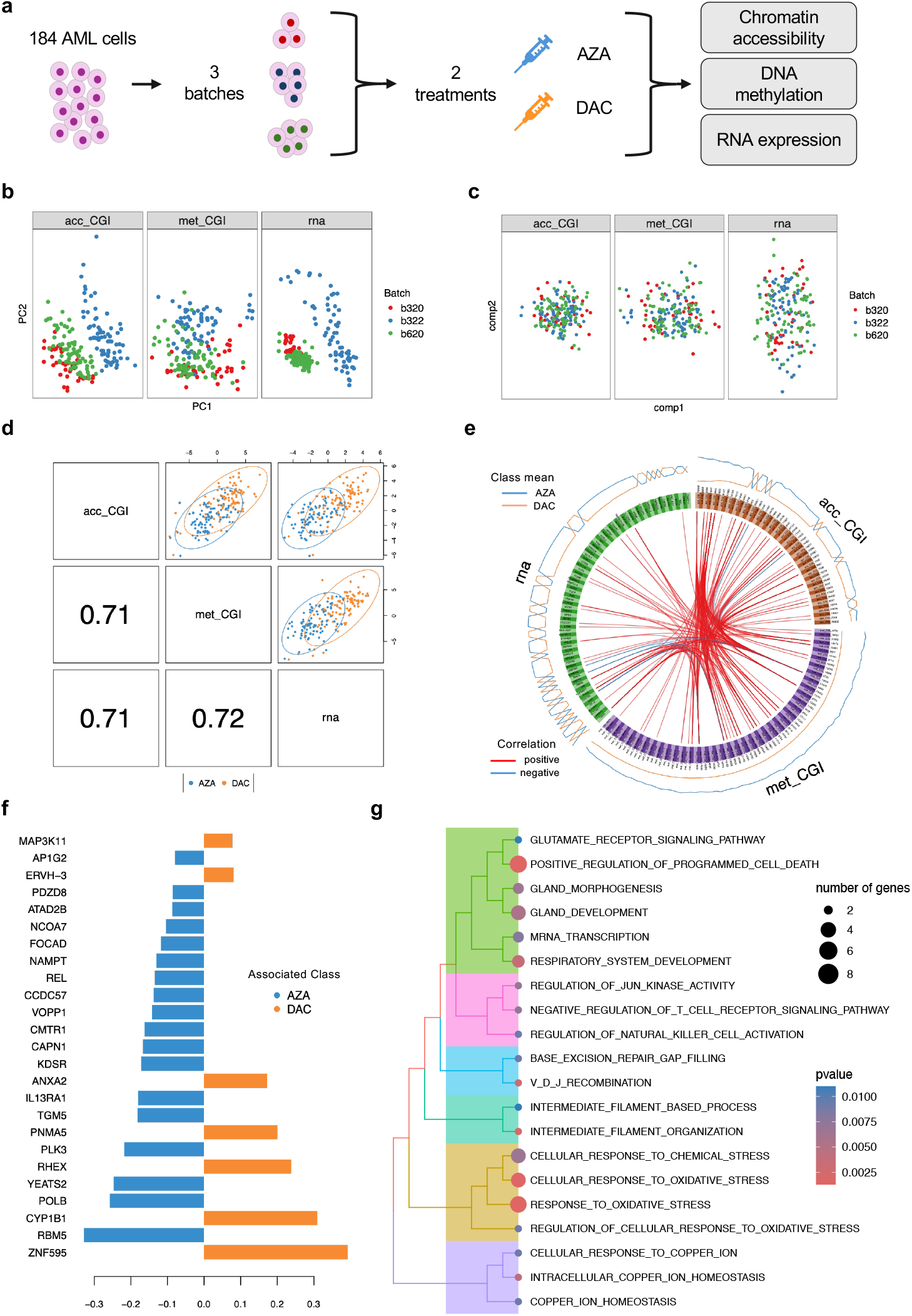
AML analysis with MINT Block. MINT Block reduces batch-driven structure and identifies a treatment-discriminating multi-omics signature in AML cells. **a)** Overview of the case study. **b)** PCA sample plots for each omics, coloured by batch. Distinct clusters show batch effects. **c)** MINT Block sample plots for each omics block, coloured by batch, showing substantially reduced batch-driven structure. **d)** Component 1 sample plots paired across omics blocks and coloured by treatment class. Pairwise correlations between component scores are shown for each pair; the components separate the treatment class and are strongly correlated across blocks. **e)** Circos plot of variables selected by MINT Block for component 1. The outer lines represent the average values observed for each treatment class and the inner lines indicate pairwise correlations between omics variables greater than 0.3 in absolute value. **f)** Loadings of RNA variables selected by MINT Block for component 1. Only the top 25 variables, ranked by absolute loading, are displayed and are coloured according to the treatment class with highest average RNA expression. **g)** GO Biological Processes over-represented among RNA variables selected by MINT Block for component 1.

#### GBR case study

For multivariate regression analysis, we used a dataset comprising 191 seawater samples collected from 48 Great Barrier Reef (GBR) sites [36]. The GBR dataset consisted of three data blocks (Figure 3a). The first block contained CLR-transformed abundances for 621 microbial taxa, providing a snapshot of the taxonomic composition of the sampled microbiomes. the second block contained CLR-transformed abundances for 4,287 functional variables derived from annotations of the genes detected in the sampled microbiomes. The third block contained 17 environmental variables aggregated by collection site, describing measurements of water chemistry, temperature and salinity. We treated this environmental data block as the response block in our analyses. The collection sites were located in seven GBR sectors, and the samples were collected across four trips conducted by the Australian Institute of Marine Science. Samples were taken from northern GBR sectors during trip 1 (November - December 2019), from southern sectors during trip 2 (January 2020), and from central sectors during trip 3 (February 2020) and trip 4 (July 2020). The sampling trips captured both spatial and temporal differences, and we treated each trip as an independent study. Pre-processing is described in Section 4.3, and sample counts by trip and reef sector are reported in Supplementary Table S2.

**Figure 3.**
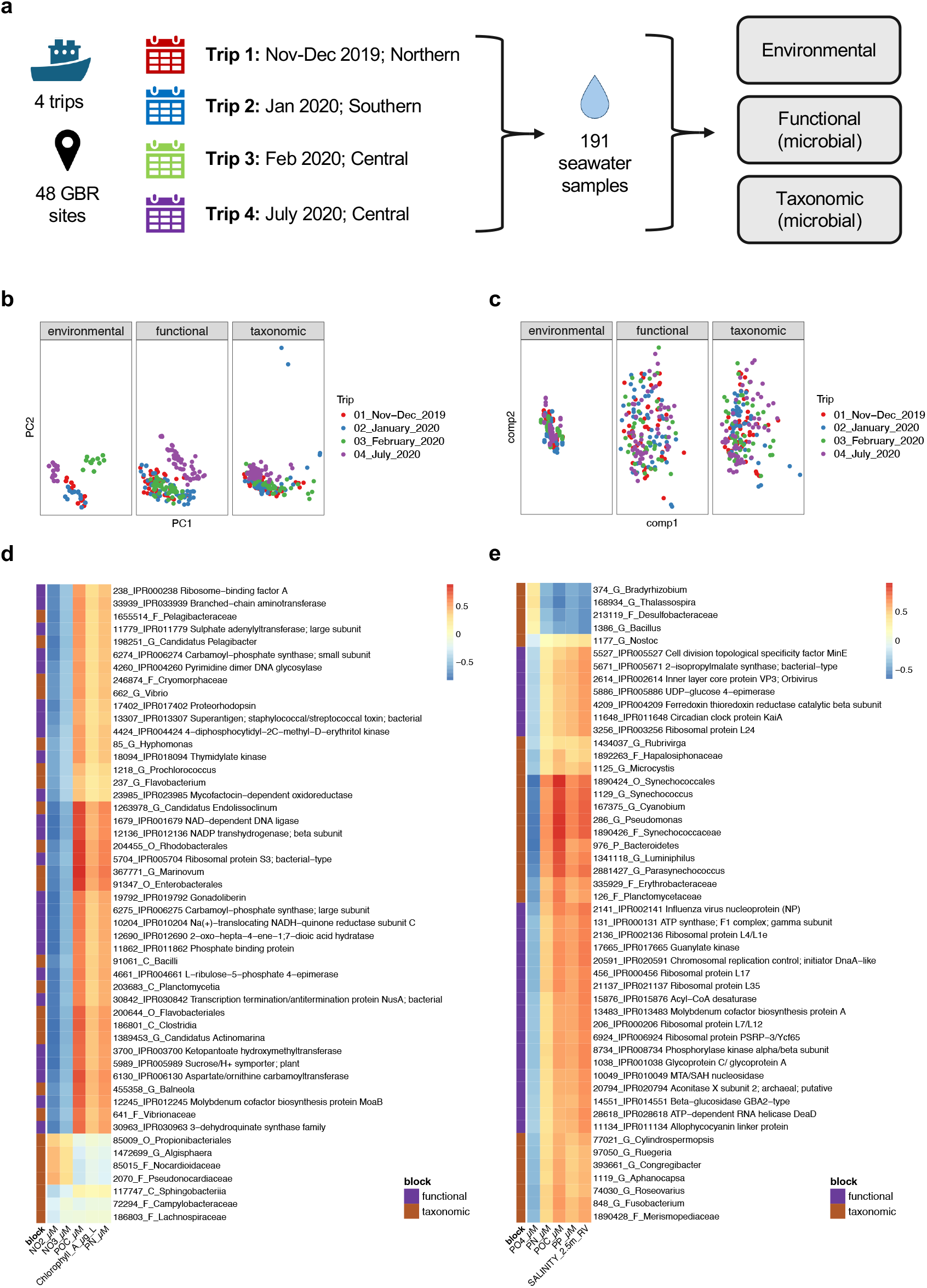
GBR analysis with MINT Block. MINT Block identifies microbial–environment relationships that are reproducible across spatio-temporally distinct GBR sampling trips.**a)** Overview of the case study. **b)** PCA sample plots for each data type, coloured by trip. Distinct clusters show strong trip-associated structure. **c)** MINT Block sample plots for each data type, coloured by trip, showing substantially reduced trip-associated structure. **d)** Heatmap of correlations between variables selected by MINT Block for component 1. **e)** Heatmap of correlations between variables selected by MINT Block for component 2. Across both components, selected microbial variables were generally negatively associated with dissolved inorganic nutrients and positively associated with particulate matter.

#### T2D case study

The R package *curatedMetagenomicData* [39] provides curated human microbiome data collected from more than 90 different studies. We combined 345 samples from five of these studies [40, 41, 42, 43, 44]; each sample was obtained from a donor with type 2 diabetes (T2D) or a healthy donor (Figure 4a). This combined dataset consisted of two microbial data blocks: the first contained abundances for 150 species, and the second contained abundances for 386 metabolic pathway variables derived using MetaCyc database annotations [45]. The T2D status of the samples served as a categorical response for classification. Section 4.4 provides details on data acquisition and pre-processing, and sample counts by status and study are reported in Supplementary Table S3.

**Figure 4.**
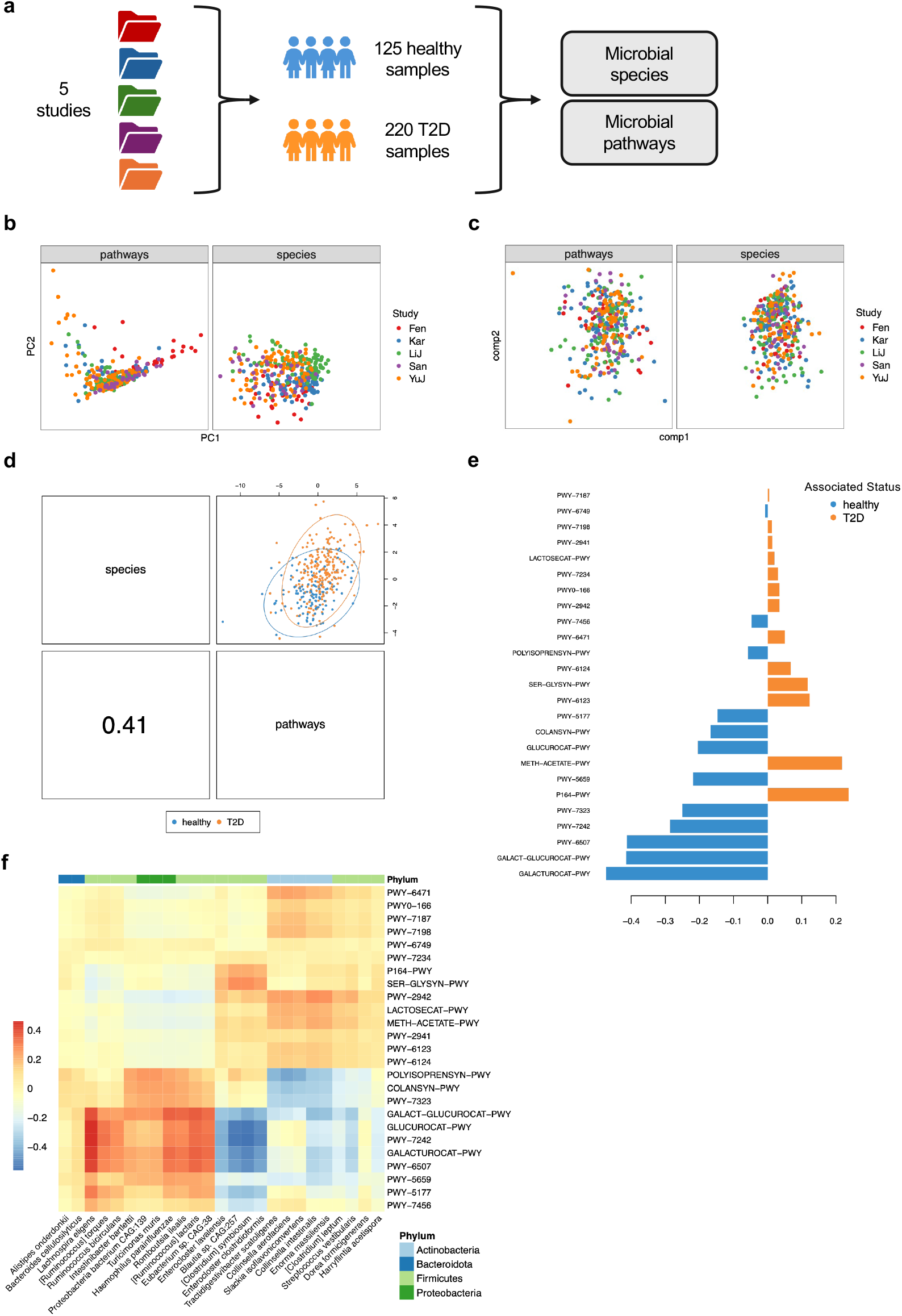
T2D analysis with MINT Block. MINT Block reduces study-associated structure and identifies linked species–pathway signatures that moderately distinguish T2D status. **a)** Overview of the case study. **b)** PCA sample plots for each data block, coloured by study. Clusters show study-associated structure. **c)** MINT Block sample plots for each data block, coloured by study. Clusters show reduced study-associated structure. **d)** Component 1 sample plots paired across data blocks and coloured by T2D status. The between-block correlation is shown; component 1 moderately separates T2D and healthy samples. **e)** Loadings of pathways selected by MINT Block for component 1, coloured according to the T2D status with the highest average pathway abundance. **f)** Heatmap of correlations between variables selected by MINT Block for component 1. Only the top 25 variables per block, ranked by absolute loading, are displayed. MINT Block identified associations between microbial species abundances and metabolic pathways abundances.

### 2.3 Overview of analyses

In Section 2.4 we benchmark the performance of MINT Block against the *mixomics* N-integration framework, which uses the same underlying PLS algorithm but does not explicitly account for study membership (Section 4.1.1). We also considered including random forests [46] and MOFA [47], but neither met all our selection criteria, which required methods to perform supervised learning and variable selection for multi-omics data. To evaluate how different approaches account for unwanted variation, we fitted both MINT Block and N-integration models with or without data correction using the batch correction methods ComBat [24] and removeBatchEffect (RBE) [31] (Section 4.5). We selected ComBat and RBE because they do not require negative control variables, are applicable to multiple types of omics, and are not restricted to settings in which relevant sample classifications are known.

We compared MINT Block with Block sPLS [48] for regression, and with DIABLO (Block sPLS-DA) [13] for classification. We evaluated the resulting lower-dimensional representations based on how effectively each approach accounted for unwanted variation (Section 4.6). To complement our evaluation, we performed Leave-One-Group-Out-Cross-Validation (LOGOCV), in which each study was held out in turn, and assessed variable selection stability [49] and prediction accuracy (Section 4.6).

In Sections 2.5–2.7, we interpret MINT Block model outputs to assess their biological relevance and illustrate how the framework can be applied across different biological contexts.

### 2.4 Benchmarking results

We first fitted models for each case study using all available samples, and calculated Adjusted Rand Index (ARI) values to measure concordance between clusters derived from the component values and study membership (Table 1, Section 4.6). Effective mitigation of study-specific variation should yield a low ARI, indicating that the clusters are not determined by study membership. Uncorrected N-integration resulted in the highest ARI values across all three case studies, indicating the need to mitigate unwanted variation. The degree to which unwanted variation affected the components varied across blocks. For example, in the AML data the highest ARI was 0.60 for the RNA block, compared to 0.38 for the Methylation block. For nearly all dataset blocks, we observed ARI values ≤ 0.05 for MINT Block models and N-integration models fitted to ComBat and RBE corrected data, indicating little residual alignment between the components study membership. The exception was N-integration + RBE for the GBR environmental block (ARI = 0.11). Therefore, by this metric, MINT Block accounted for unwanted variation during model fitting as effectively as applying prior data correction with an established method.

**Table 1.** Benchmarking results. N-integration models were fitted to uncorrected data and to data corrected using ComBat or removeBatchEffect (RBE), whereas MINT Block was fitted directly to uncorrected data. Performance was evaluated using the adjusted Rand index (ARI) for alignment between component-based clusters and study membership, variable-selection stability, balanced error rate (BER) for classification, and root mean squared error (RMSE) for regression. Lower ARI, BER and RMSE values and higher stability values indicate better performance. Bold font indicates the best result for each criterion within a dataset or data block. MINT Block achieved prediction errors lower than those of uncorrected and corrected N-integration models while effectively accounting for study-specific variation.

| Score | Dataset |  | N-integration |  |  | MINT Block |
| --- | --- | --- | --- | --- | --- | --- |
|  |  |  | Uncorrected | ComBat | RBE |  |
| ARI | AML | Accessibility | 0.51 | **0.01** | 0.02 | 0.02 |
|  |  | Methylation | 0.38 | 0.06 | 0.03 | **0.02** |
|  |  | RNA | 0.60 | 0.04 | **0.00** | 0.04 |
|  | GBR | Taxonomic | 0.31 | 0.05 | **0.02** | 0.04 |
|  |  | Functional | 0.27 | **0.03** | 0.05 | **0.03** |
|  |  | Environmental | 0.55 | 0.03 | 0.11 | **0.01** |
|  | T2D | Species | 0.13 | 0.01 | 0.01 | **0.00** |
|  |  | Pathways | 0.17 | 0.01 | 0.02 | **0.00** |
| Stability | AML | Accessibility | 0.38 | 0.42 | 0.41 | **0.44** |
|  |  | Methylation | 0.38 | **0.53** | 0.49 | 0.52 |
|  |  | RNA | **0.41** | 0.38 | 0.38 | 0.38 |
|  | GBR | Taxonomic | 0.49 | **0.57** | 0.55 | **0.57** |
|  |  | Functional | 0.33 | **0.45** | 0.39 | 0.43 |
|  | T2D | Species | 0.61 | **0.62** | 0.58 | **0.62** |
|  |  | Pathways | 0.43 | **0.44** | **0.44** | 0.43 |
| BER | AML | Mean | 0.41 | 0.33 | 0.36 | **0.24** |
|  |  | SD | 0.14 | 0.17 | 0.20 | 0.17 |
|  | T2D | Mean | 0.42 | 0.41 | 0.41 | **0.35** |
|  |  | SD | 0.07 | 0.03 | 0.04 | 0.06 |
| RMSE | GBR | Mean | 1.30 | 1.24 | 1.27 | **1.22** |
|  |  | SD | 0.09 | 0.06 | 0.08 | 0.05 |

We next performed LOGOCV analysis for each dataset and compared variable selection stability scores and prediction errors (described in Section 4.6). Overall, addressing unwanted variation resulted in higher stability scores, indicating more consistent variable selection across different subsets of studies or batches. Variables selected consistently across these subsets may represent more generalisable biomarker signatures (Table 1). Removing unwanted variation with ComBat led to the highest stability scores overall, followed closely by MINT Block. During LOGOCV, we assessed the predicted responses for held-out observations using either balanced error rate (BER) for classification or root mean squared error (RMSE) for regression (Table 1, Supplementary Figures S1-S2). BER gives equal weight to each class and is therefore appropriate to imbalanced class designs. MINT Block achieved the lowest BER for both the AML (0.24) and T2D (0.35) datasets, and the lowest RMSE for the GBR dataset (1.22). Thus, although ComBat generally produced the most stable variable selection, MINT Block achieved the best predictive performance across all three datasets, suggesting that its lower-dimensional representations retained more response-relevant signal.

### 2.5 AML case study

To visualise unwanted variation in the AML data, we performed principal component analysis (PCA) separately for each omics block (Figure 2b). Across all omics blocks, PCA plots showed that samples clustered by batch rather than treatment class. The batch effect was especially strong in the Accessibility and RNA data, consistent with the ARI values obtained from our benchmarking for uncorrected N-integration (Table 1). To account for unwanted variation, we fitted a two-component MINT Block classification model with treatment class as the response. Each component selected 50 Accessibility, 75 Methylation, and 75 RNA variables (Section 4.5). The resulting sample plots differed from the PCA plots and showed no clear clustering by batch, consistent with substantially reduced batch-driven structure in the MINT Block lower-dimensional representation (Figure 2c).

Our tuning indicated that a model with a single component was optimal (Section 4.5). We therefore focused our interpretation on component 1, which separated cells according to treatment class. The pairwise correlations between component scores across block pairs were all ≥ 0.70, indicating that MINT Block captured a latent shared signal underlying all omics blocks (Figure 2d). Most selected variables were positively correlated and had higher average values in the AZA treatment group than in the DAC group (Figure 2e).

To assess the biological relevance of the fitted model, we examined the top 25 RNA variables ranked by the magnitude of their component 1 loading (Figure 2f). Eighteen had higher average expression in the AZA group and seven in the DAC group. Among the AZA-associated genes, RBM5, NAMPT and NCOA7 have reported roles in AML cell survival [50, 51, 52]. Among the DAC-associated genes, CYP1B1 has been linked to AML progression and RHEX has been proposed as a therapeutic target [53, 54, 55]. GO over-representation analysis of all selected RNA variables highlighted processes related to cell death, oxidative stress, immune signalling, DNA repair, intermediate-filament organisation and copper homeostasis (Figure 2g). Detailed gene-level evidence and the complete interpretation of the enriched processes are provided in Supplementary Section S1.1 and Supplementary Table S4.

In summary, MINT Block accounted for pronounced batch effects and identified genes with differing average expression between AZA and DAC. These genes and their associated biological processes have reported links to AML and cancer. Together, the findings generate hypotheses about the treatments’ distinct molecular effects in AML.

### 2.6 GBR case study

PCA plots of each GBR data block revealed strong clustering by sampling trip, particularly between summer and winter samples (Figure 3b). Because each reef was sampled only once across different regions and seasons, these dominant spatio-temporal effects could mask finer-scale microbial–environment relationships. Our NP-integration analysis therefore aimed to identify correlated taxonomic, functional and environmental signatures and assess whether they were reproducible across the independent sampling trips.

We fitted a two-component MINT Block multivariate regression model using the environmental data as the response. Each component selected 25 taxonomic variables, 25 functional variables, and 5 environmental variables (Section 4.5). The MINT Block sample plots showed little residual clustering by trip, suggesting that the model identified coordinated taxonomic, functional, and environmental variables that were reproducible across independent sampling trips (Figure 3c).

Nearly all taxonomic and functional variables selected for MINT Block component 1 were negatively correlated with nitrite (NO_2_) and nitrate (NO_3_) concentrations, and positively correlated with particulate organic carbon (POC), chlorophyll *a*, and particulate nitrogen (PN) (Figure 3d, see Section 4.6 for how correlations were calculated). Microbial variables selected for MINT Block component 2 showed a similar pattern: they were negatively correlated with dissolved inorganic nutrients including phosphate (PO_4_) concentrations, and positively correlated with particulates (PN, POC, and particulate phosphorus - PP) as well as salinity (Figure 3e).

The taxa most strongly associated with higher particulate organic matter and lower dissolved inorganic nitrogen included picocyanobacteria from the order *Synechococcales*, particularly *Synechococcus, Cyanobium* and *Parasynechococcus*. This pattern is consistent with the proposed use of the *Synechococcus*:*Prochlorococcus* abundance ratio as an indicator of nutrient enrichment on coral reefs [56]. Selected heterotrophic taxa included members of *Flavobacteriales* and *Rhodobacterales*, which can metabolise phytoplankton-derived organic substrates [57]. The selected functional variables represented core metabolism, respiration, ATP generation and biosynthetic activity. Together, these variables support a coordinated microbial response involving nutrient drawdown, biological production and organic-matter utilisation. A more detailed ecological interpretation is provided in Supplementary Section S1.2.

Overall, MINT Block integration recovered microbial–environmental relationships that were reproducible across spatio-temporally distinct sampling trips, linking microbial signatures to lower dissolved nutrient concentrations and higher particulate matter.

### 2.7 T2D case study

For both the species and pathways blocks, PCA plots showed moderate clustering by study (Figure 4b). This study-associated structure was substantially reduced from the two-component MINT Block classification model (Figure 4c). Component 1 selected 50 species variables and 25 pathways variables, while component 2 selected 75 from each block (Section 4.5).

MINT Block component 1 showed partial separation by T2D status and a moderate between-block correlation of 0.41 (Figure 4d). In contrast, component 2 was only minimally able to discriminate T2D status but was more highly correlated between blocks (Supplementary Figure S3). We therefore focussed on component 1, to examine the microbial variables associated with T2D.

Among the pathway variables selected for component 1, the strongest contributors to discrimination had higher average abundance in healthy samples (Figure 4e). These included pathways involved in colanic acid and GDP-mannose biosynthesis and the degradation of *β*D-glucuronide and D-glucuronate, which have previously been reported as diabetes-associated biomarkers [58, 59, 60]. Pathways with higher average abundance in T2D samples included purine and pyrimidine metabolism, methanogenesis and amino-acid biosynthesis; purine metabolism has previously been associated with diabetes progression [61]. Detailed pathway-level evidence, including associations whose biological interpretation remains uncertain, is provided in Supplementary Section S1.3 and Supplementary Tables S5 and S6.

Correlations between the top 25 species and pathway variables selected for component 1 identified three groups of species (Figure 4f). Health-associated species, including *Lachnospira eligens* and several *Ruminococcus* species, were positively correlated with pathways that were also more abundant in healthy samples. Two groups composed predominantly of T2D-enriched species, including *Enterocloster lavalensis, Collinsella aerofaciens* and *Streptococcus vestibularis*, were correlated with pathways involving purine or pyrimidine metabolism, methanogenesis and amino-acid biosynthesis. The complete species groups and supporting pathway associations are detailed in Supplementary Section S1.3.

In summary, MINT Block reduced study-associated structure despite only moderate separation by T2D status. It identified cross-study associations between microbial species and metabolic pathway abundances with reported relevance to T2D and metabolic health.

## 3 Discussion

We presented the MINT Block NP-integration framework, which addresses the need for methods that can integrate multi-omics data from independent studies. MINT Block performs dimensionality reduction and variable selection within multivariate regression or classification analysis, while simultaneously accounting for study-specific variation arising when multiple studies are combined.

Using data from several biological contexts, we benchmarked MINT Block against the conventional sequential NP-integration workflow, in which each omics blocks is corrected independently before the application of an N-integration method. This benchmarking demonstrated that MINT Block identified stable multi-omics signatures and achieved higher predictive accuracy than the sequential approaches, without requiring separate data-correction steps. Detailed analysis of the three case studies provided insights into genes and biological processes associated with AZA and DAC treatment in acute myeloid leukemia, reproducible microbial-environment relationships across the Great Barrier Reef, and species-pathway abundance patterns associated with type 2 diabetes.

When fitting a MINT Block model, it is necessary to specify the number of components and the number of variables to select for each component. To this end, we tuned the models using cross-validation to determine which configurations provided the best predictive performance. We primarily used existing *mixOmics* tuning procedures, modified for NP-integration. The exception was our novel use of the covariance ratio for variable selection in the regression context. Although our tuning methods have the appealing feature of emphasising generalisability to new observations, prediction-focused tuning is not always optimal for addressing the biological questions driving an analysis. In the acute myeloid leukemia analysis, for example, the number of RNA variables selected based on tuning was too large to describe concisely, yet was barely sufficient for testing for over-representation of GO terms. Likewise, in the Great Barrier Reef analysis, we found it advantageous to distribute the tuned number of variables across two components, despite tuning indicating that a single component was optimal. In addition, cross-validation is time consuming, which is a barrier to evaluating all possible variable selection configurations. We thus advise MINT Block users to treat the results of tuning as a guide, and to consider their research objectives when selecting the number of components and variables. Accordingly, future work could explore other methods for determining the number of components and variables, including alternative optimisation criteria such as explained variance [62], or statistical tests on loadings to remove the need for cross-validation [63].

Another important consideration is whether to integrate all available studies or use only a subset. When creating the type 2 diabetes dataset, we excluded one study because the data included repeated measurements from a small number of donors. This highlighted the importance of study-level quality control during pre-processing. Even after curation, the predictive performance of models fitted to the combined dataset remained low, which may reflect either a limit to the amount of heterogeneity that MINT Block can accommodate or a weak disease-related signal shared across studies. Future work should therefore evaluate how MINT Block model performance changes with the number of included studies and the degree of within- and between-study heterogeneity, with the aim of developing guidelines for users. In settings where careful study selection still produces a poor model fit, incorporating prior biological knowledge from molecular interaction databases into MINT Block may improve its robustness to high levels of noise [16], as illustrated by the dimensionality reduction methods f-scLVM [64] and PLIER [65].

Further extensions are motivated by limitations of the data we used in this study. Our evaluation was restricted to used bulk multi-omics datasets and one small single-cell multi-omics dataset, all containing real data. We are not aware of a simulation tool that can simultaneously reproduce (i) complex, non-linear relationships between molecules and (ii) realistic study-specific variation while preserving the known ground truth required to evaluate variable selection in supervised analyses. Developing such a tool would enable more comprehensive benchmarking of multi-omics integration methods.

In addition, the increasing prevalence of large-scale, single-cell multi-omics data creates a need to better understand the scalability of MINT Block, and may require addressing the computational complexity often associated with single-cell data [66]. Addressing this additional complexity could involve either employing a pseudo-bulk approach [67], or further developing the MINT Block integration algorithm.

In conclusion, to our knowledge, MINT Block is the first omics analysis tool to simultaneously integrate multiple types of omics data from independent studies. Given the increasing availability of multi-omics datasets, MINT Block provides researchers with a unified approach to combine studies and derive comprehensive, reproducible multi-omics signatures.

## 4 Methods

### 4.1 The MINT Block NP-integration framework

Consider a multi-block dataset *X* formed by combining *M* ≥ 2 independent studies, each contributing the same *Q* ≥ 3 data blocks: *Q* − 1 explanatory blocks and one response block (Figure 1). Each block contains a different set of variables measured on the same 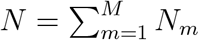 observations, where *N*_m_ is the number of observations from study *m*. For block *q* we denote the data matrix of size (*N* × *P*_q_) by *X*^(q)^, where *P*_q_ is the number of variables. We partition the data by study and denote the matrix of size (*N*_m_ × *P*_q_) for study *m* and block *q* by *X*^(m,q)^.

MINT Block estimates *H* latent components and *H* corresponding loading vectors for each block, yielding a lower-dimensional representation of *X*. For each component *h* ∈ {1, …, *H*}, the covariance among the block-specific components is jointly maximised and informative variables can be selected by imposing sparsity on the loading vectors.

MINT Block is formulated for supervised analysis, in which the *Q*th block contains one or more response variables and blocks *q* ∈ {1, …, *Q* − 1} contain distinct sets of explanatory variables. We denote the response matrix by *Y* = *X*^(Q)^ (consistent with [13] for N-integration). MINT Block performs multivariate regression for quantitative response variables. For classification, *Y* is an indicator matrix obtained by dummy coding a categorical variable representing class membership.

The framework consists of four methods: MINT Block PLS (multivariate regression without variable selection), MINT Block sPLS (multivariate regression with variable selection), MINT Block PLS-DA (classification without variable selection), and MINT Block sPLS-DA (classification with variable selection). A common algorithm underlies all four methods and combines the N-integration and P-integration frameworks implemented in the *mixOmics* [48] R package as described below (Section 4.1.3).

Throughout this section, for centred vectors *u* and *v* of length *n*, we define

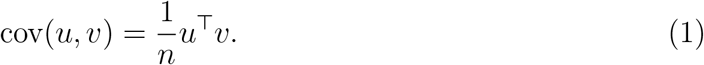

#### 4.1.1 The N-integration framework

N-integration in *mixOmics* [6] performs multivariate regression and classification analyses for datasets consisting of *Q* ≥ 3 blocks of data. These methods take column-centred and scaled matrices *X*^(q)^, *q* ∈ {1, …, *Q*}, as input and estimate *H* block-specific latent components and their associated loading vectors. For each component *h* ∈ {1, …, *H*}, the N-integration optimisation problem is:

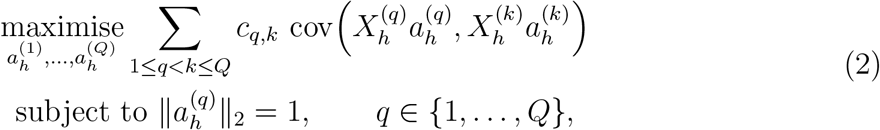

with the additional constraint

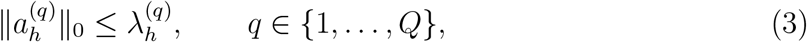

for the sparse variants.

Here, 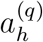 is the loading vector of length *P*_q_ for block *q* and component *h*; 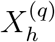 is the corresponding residual matrix (Algorithm 1); and

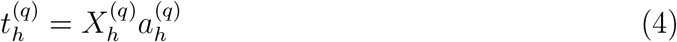

is the component score vector. The coefficient *c*_q,k_ is the (*q, k*)th element of a symmetric design matrix *C*, with *c*_q,q_ = 0, and specifies the weight assigned to the covariance between blocks *q* and *k*.

For the sparse variants, 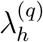 specifies the maximum number of non-zero elements in 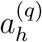. Variables with non-zero loading coefficients are selected for block *q* and component *h*. Sparsity is imposed on an unthresholded loading update *ã* using the following soft-thresholding operation:

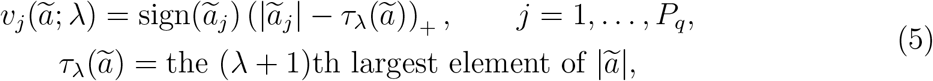

where (*x*)_+_ = max(*x*, 0) and *τ*_λ_(*ã*) = 0 when *λ* = *P*_q_. The thresholded loading vector is then normalised as

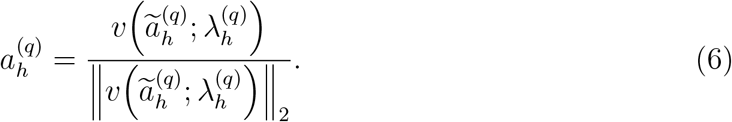

N-integration methods compute loading vectors and components sequentially. They first estimate 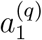 for 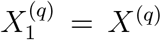, *q* ∈ {1, …, *Q*}, by solving the optimisation problem in equation (2). For each successive component, the loadings and component scores are computed from new residual matrices obtained through deflation. For explanatory block *q < Q*, the residual matrix for the next component is

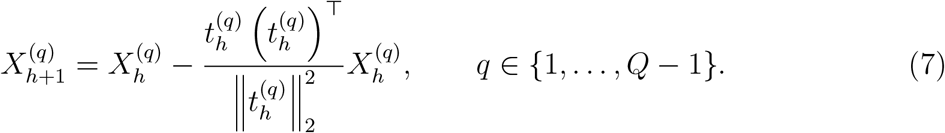

The response block is deflated using the average projection across the *Q* − 1 explanatory components:

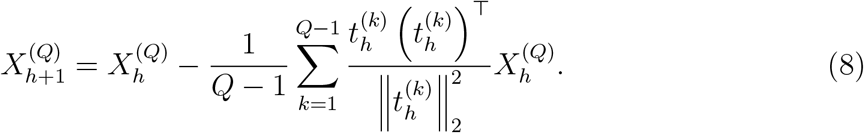

Algorithm 1 summarises how these equations are applied across the blocks.

##### Algorithm 1

Deflation

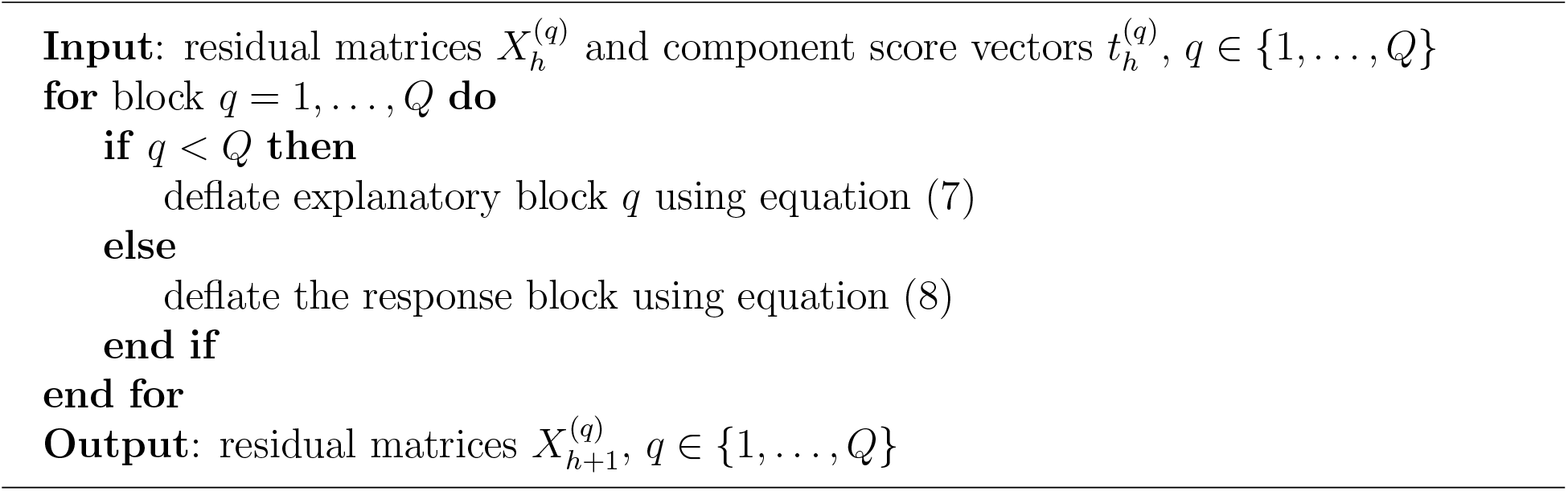

The *mixOmics* N-integration methods we use in this study are Block sPLS [6] for regression with variable selection, and Block sPLS-DA (also known as DIABLO; [13]) for classification with variable selection.

#### 4.1.2 The P-integration framework

The *mixOmics* P-integration framework [22] comprises methods for multivariate regression and classification of two data blocks measured across multiple studies. For each study *m*, the first block contains explanatory variables and the second contains the response variables, such that *X*^(m,2)^ = *Y* ^(m)^. Each study contains the same variables within a given block.

P-integration methods take column-centred and scaled matrices *X*^(m,q)^, *m* ∈ {1, …, *M*} and *q* ∈ {1, 2}, as input. The centring and scaling are performed within each study. The methods estimate *H* study-specific component score vectors and corresponding global loading vectors shared across studies.

For each component *h* ∈ {1, …, *H*}, the P-integration optimisation problem is:

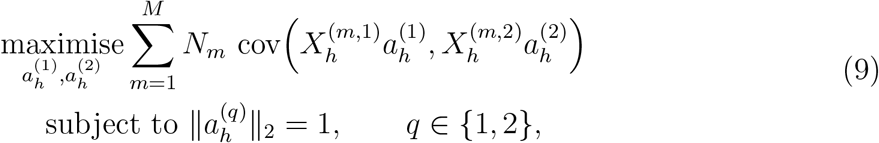

with the additional constraint

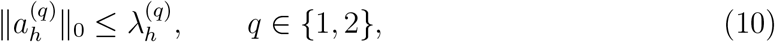

for the sparse variants.

Here, 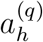 is the global loading vector for block *q* and component *h*, shared across all studies; 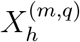 is the residual matrix for study *m* and block *q*; and

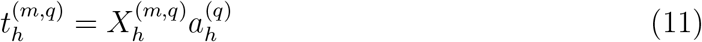

is the corresponding study-specific component score vector. The factor *N*_m_ weights the contribution of each study according to its sample size. For sparse methods, 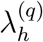 specifies the maximum number of non-zero elements in 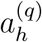.

P-integration methods solve the optimisation problem in equation (9) by computing study-specific (*partial*) loading vectors 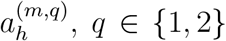 and *m* ∈ {1, …, *M*}. These partial loading vectors are aggregated across studies to obtain the global loading vectors 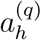, *q* ∈ {1, 2}. For the sparse variants, soft thresholding is applied to the aggregated loading vectors before they are normalised. The P-integration methods implemented in *mixOmics* are MINT PLS for multivariate regression without variable selection, MINT sPLS for multivariate regression with variable selection, MINT PLS-DA for classification without variable selection, and MINT sPLS-DA for classification with variable selection.

#### 4.1.3 The MINT Block NP-integration algorithm

MINT Block combines the N-integration and P-integration frameworks. For each component *h* ∈ {1, …, *H*}, it solves the following optimisation problem:

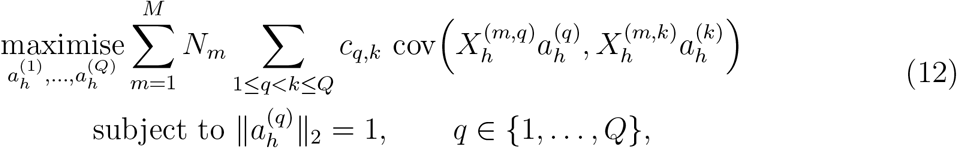

with the additional constraint

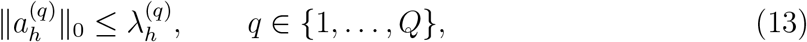

for the sparse variants.

Here, 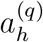 is the global loading vector of length *P*_q_ for block *q* and component *h*, shared across studies; 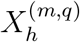 is the residual matrix for study *m* and block *q*; and

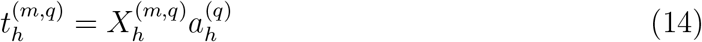

is the corresponding study-specific component score vector. The coefficient *c*_q,k_ is defined as in equation (2). Under the covariance definition in equation (1), the factor *N*_m_ weights each study by its sample size and yields the cross-product updates used in Algorithm 2.

We solve equation (12) using Algorithm 2:

##### Algorithm 2

MINT Block NP-integration algorithm

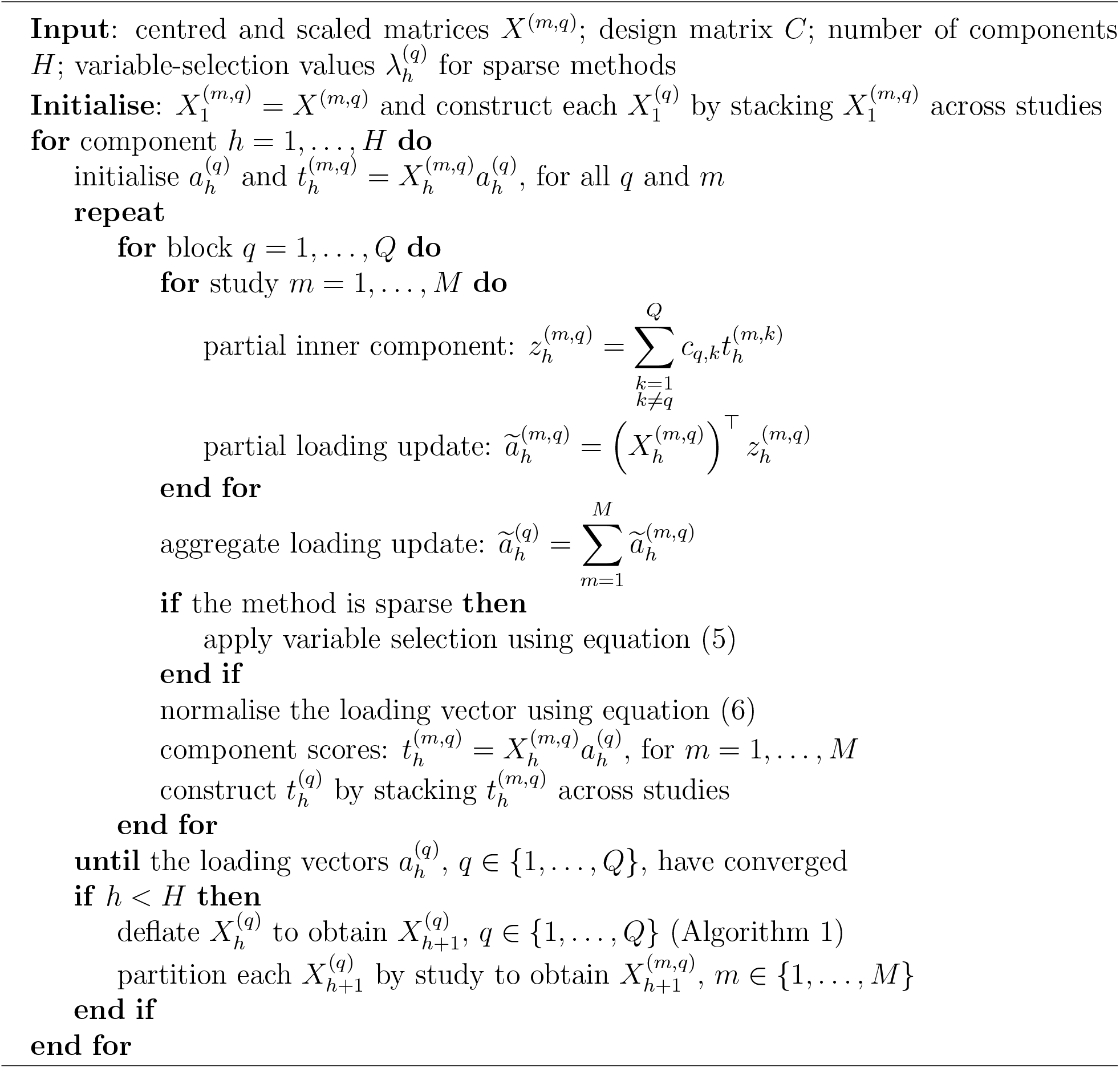

After all *H* components have been estimated, the algorithm returns the component matrix

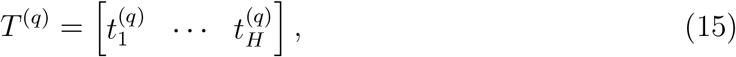

of size (*N* × *H*), and the loading matrix

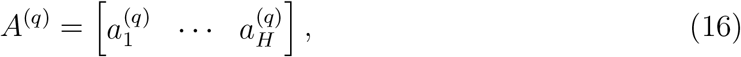

of size (*P*_q_ ×*H*), for each block *q*. The component matrices provide lower-dimensional representations of the observations, while the loading matrices contain the variable weights. For sparse methods, variables with non-zero loading coefficients are selected.

#### 4.1.4 Prediction for new data

A fitted MINT Block model uses its component and loading matrices to predict responses for new observations, following the prediction procedure used by *mixOmics* N-integration models [48, 13].

Let *X*^(new,q)^, *q* ∈ {1, …, *Q* − 1}, denote the explanatory-block matrices for *N*_new_ new observations. Each matrix is of size (*N*_new_ × *P*_q_) and must contain the same variables, in the same order, as the corresponding training block. If the observations are from multiple studies, the data from each study are centred and scaled independently using the same preprocessing procedure applied during model fitting.

The objective is to estimate 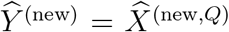, the matrix of predicted response values. We first obtain a block-specific prediction from each explanatory block:

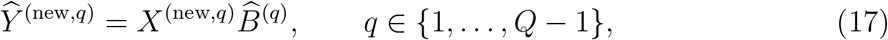

where *Ŷ* ^(new,q)^ contains the response predictions obtained from explanatory block *q*, and 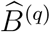 is the corresponding regression-coefficient matrix:

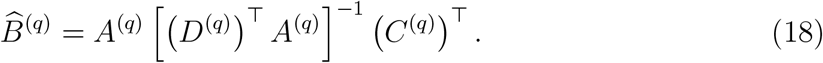

Here, *A*^(q)^ is the loading matrix of size (*P*_q_ ×*H*) for explanatory block *q*. The matrices *D*^(q)^ and *C*^(q)^ are the transposes of the least-squares coefficient matrices obtained by regressing *X*^(q)^ and *Y*, respectively, on the component matrix *T* ^(q)^. Equivalently, *T* ^(q)^ (*D*^(q)^) ^⊤^ and *T* ^(q)^ (*C*^(q)^)^⊤^ are the orthogonal projections of *X*^(q)^ and *Y* onto the column space of *T* ^(q)^. These loading matrices are calculated as

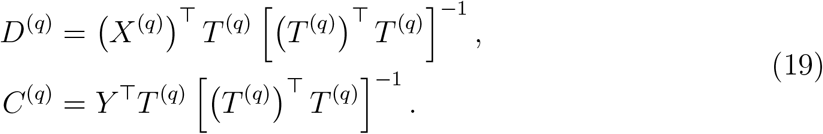

We then use a simple or weighted average of the block-specific predictions to calculate the consensus prediction *Ŷ* ^(new)^.

In addition to predicting response values, we predict component scores for new observations as

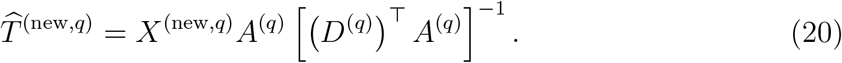

For classification, the predicted response or component scores can be converted into class predictions using several decision rules, as described in [48]. Under the *maximum distance* rule, observation *i* is assigned to the class with the largest predicted indicator value:

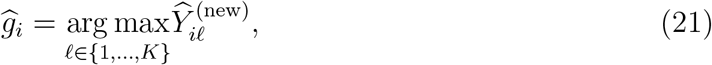

where *K* is the number of classes. Alternatively, class predictions can be obtained separately from each *Ŷ* ^(new,q)^ and combined using majority or weighted voting. Class predictions can also be based on centroid or Mahalanobis distances between the rows of 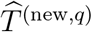 and class centres estimated from the training component scores *T* ^(q)^. The resulting block-specific class predictions are then combined using majority or weighted voting.

#### 4.1.5 Tuning the number of components

We tune the number of components in MINT Block models using prediction performance assessed by LOGOCV.

Given a maximum number of components 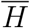, we perform LOGOCV (Section 4.6) for each 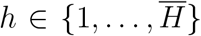, fitting MINT Block PLS(-DA) models with *h* components on the LOGOCV training sets and evaluating *predicted* values across the left-out folds (Section 4.1.4). We use different selection criteria for multivariate regression and classification:

##### Multivariate regression

In addition to LOGOCV, for each 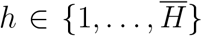 we fit a MINT Block PLS model using all studies, with no held-out folds, to obtain *fitted* values for the response variables. We then select the optimal number of components based on *Q*^2^ values [11, 68], calculated as

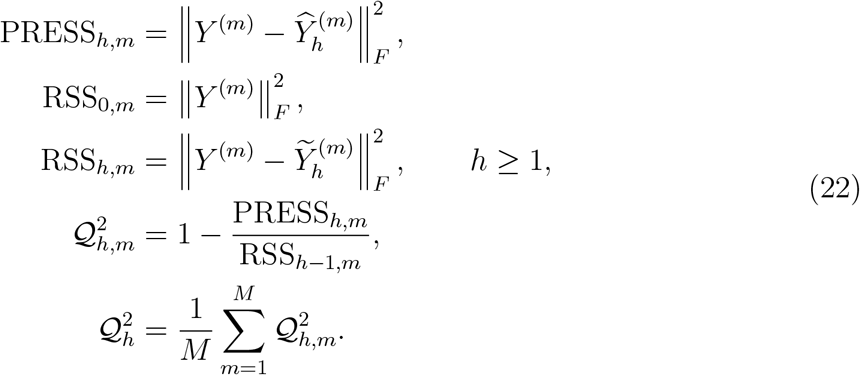

Here, ∥*A*∥_F_ is the Frobenius norm of matrix *A*; *Y* ^(m)^ is the matrix of observed response values for study 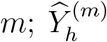 is the matrix of response values predicted for study *m* during LOGOCV using a model with *h* components; and 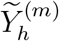 is the matrix of fitted response values for study *m* from the model with *h* components fitted using all studies. Because *Y* ^(m)^ is centred and scaled within study, RSS_0,m_ = (*N*_m_−1)*P*_Q_ when each response variable has unit sample variance.

We select *H* using a threshold of 0.0975. If 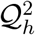 first falls below this threshold at component *h*^⋆^ *>* 1, we select *H* = *h*^⋆^ − 1. If 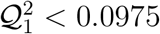, we retain one component as the minimum required to fit the model but consider the model to have limited predictive ability. If no candidate value falls below the threshold, we select 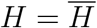.

##### Classification

We select the value of 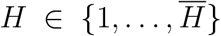 that yields the lowest mean balanced error rate (BER) across the held-out studies (Section 4.6).

#### 4.1.6 Tuning the number of variables

After selecting the number of components, we tune the values of 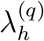, which determine the maximum number of variables selected per block and component (Equation 13). For each block *q* ∈ {1, …, *Q*}, we specify a set of candidate values for 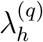. We tune the components sequentially. Before tuning component *h*, we fix the variable-selection parameters for all preceding components *g* ∈ {1, …, *h* − 1} at their previously selected values. We then evaluate the candidate combinations 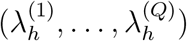 for component *h*. We tune these parameters using LOGOCV (Section 4.6) and evaluate predictions for the held-out studies (Section 4.1.4). We use different selection criteria for multivariate regression and classification:

##### Multivariate regression

For each *h* ∈ {1, …, *H*}, we fit MINT Block sPLS models using all studies in addition to performing LOGOCV. For the current component *h*, we fit a separate model for each candidate combination of 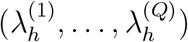, while holding the parameters for preceding components fixed. For each candidate combination, we calculate the covariance ratio:

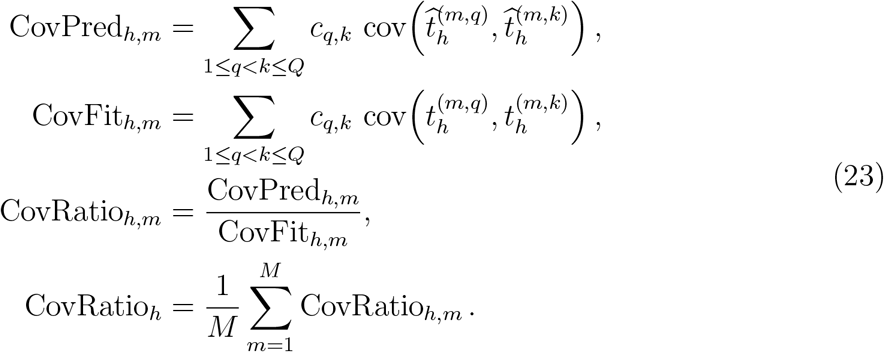

Here, *c*_q,k_ specifies the relative importance of the connection between blocks *q* and *k* (Section 4.1); 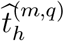is the out-of-sample component score vector for held-out study *m*, component *h*, and block *q*; and 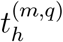 is the corresponding fitted component score vector from the model trained using all studies. The covariance ratio is defined when CovFit_h,m_ ≠0. We select the combination 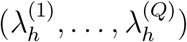 that yields the highest CovRatio_h_ for component *h*.

##### Classification setting

We set 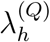 such that all response variables are selected. We then select the combination 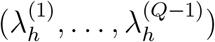 that yields the lowest mean BER across the held-out studies (Section 4.6) for component *h*.

### 4.2 AML case study data acquisition and pre-processing

We analysed a subset of the single-cell multi-omics dataset from [38]. The dataset had been generated by profiling acute myeloid leukemia (AML) cells using scNMT-seq, a method that profiles DNA methylation, chromatin accessibility, and RNA expression. For the DNA methylation and chromatin accessibility data, the variables represented CpG and GpC sites, respectively. The RNA counts had been normalised with the SCTransform method from Seurat [69], and subsequently log(*x*+1) transformed. The dataset comprised three experimental batches, which we treated as three independent studies in our analysis.

To create the analysis subset, we selected cells that had been treated with either azacytidine (AZA) or decitabine (DAC). We then excluded RNA variables lacking an Ensembl ID or having a normalised count of ≥ 5 in fewer than 10% of cells. We used the log(*x* + 1) transformed normalised counts of the retained RNA variables in the analysis. For the DNA methylation and chromatin accessibility data, we retained only sites located within CpG islands (CGIs), and excluded variables for which more than 40% of cells had a missing value. For each data type, we retained up to the 5,000 variables with the highest variance across all studies. We then excluded variables with no variance in one or more studies.

### 4.3 GBR case study data acquisition and pre-processing

We obtained metagenomics and physico-chemical data from the published GBR study [36] downloaded from the Australian Institute of Marine Science (AIMS) data repository: https://apps.aims.gov.au/metadata/view/5956fd44-f864-4811-9ec3-15499f1863c4. The data had been generated from surface seawater samples (target depth of 5 m) collected from 48 offshore reefs across the Great Barrier Reef during four separate sampling trips conducted between November 2019 and July 2020, across the austral summer and winter.

In the original study, seawater samples were pre-filtered through 5-*µ*m filters to remove large debris and eukaryotic cells prior to microbial biomass collection on 0.22-*µ*m Sterivex filters for shotgun metagenomic sequencing [36]. Quality-filtered reads were aligned to the NCBI nr database using DIAMOND [70], retaining only the top hit for each read with an e-value *<* 10^−5^. Taxonomic and functional annotations were generated using MEGAN [71] and exported as taxonomic and Gene Ontology (GO) functional profiles. Where genus-level or GO-level annotations were unavailable, higher taxonomic or UniProt classifications were retained. Pre-processing steps included removing (1) non-annotated reads, (2) eukaryotic and viral annotations, (3) prokaryotic reads annotated only at the domain level, and (4) rare taxonomic or functional features with relative abundances below 0.0001%. The resulting microbial abundance data were centred log-ratio (CLR) transformed after the addition of pseudocounts to account for the compositional and sparse nature of metagenomic sequencing data [72].

For each reef site, median values of 17 physico-chemical variables—including temperature, salinity, picoplankton pigments, and particulate and dissolved nutrients—were calculated from three replicate measurements (detailed protocol in [36]).

We used these data to create a dataset consisting of three blocks: (1) a microbial taxonomy block containing abundances for 621 taxonomic variables, (2) a microbial functional block containing abundances for 4,287 functional variables based on gene annotations, and (3) an environmental block containing measurements for 17 physico-chemical variables aggregated by reef site. The taxonomic and functional blocks were CLR transformed. Because each reef site was sampled only once, the effects of geography and sampling time were confounded. We therefore treated each sampling trip as an independent study in our analysis.

### 4.4 T2D case study data acquisition and pre-processing

The *curatedMetagenomicData* R package provides uniformly processed human microbiome data generated from metagenomic sequencing [39]. The repository contains multiple types of microbiome data for *>* 22,000 samples, covering more than 30 diseases and *>* 90 studies. We created a dataset by combining data from five different studies [40, 41, 42, 43, 44], selecting samples from donors with T2D or healthy donors. The resulting dataset contained two types of microbial data: (1) species-level taxonomic profiles in the form of relative abundances and (2) metabolic pathway abundances.

Within each combination of data type, study, and participant status (T2D or healthy), we removed variables observed in fewer than 10% of samples. We then applied the CLR transformation. We merged the studies and removed variables that were not observed in every study.

### 4.5 Model fitting

For the AML and T2D data, we fitted NP-integration (MINT Block sPLS-DA) and N-integration (DIABLO) [13] models, using the indicator matrix representing class membership as the response. For the GBR data, we fitted NP-integration (MINT Block sPLS) and N-integration (Block sPLS) [48] models, using the matrix of environmental data as the response. We fitted MINT Block and N-integration models to the uncorrected data. We additionally fitted N-integration models to data corrected using ComBat [24] and removeBatchEffect (RBE) [31]. Both ComBat and RBE model known extraneous factors to estimate and remove unwanted variation while preserving variation associated with specified biological covariates. RBE applies an additive linear-model adjustment, whereas ComBat uses empirical Bayes shrinkage to remove both additive and multiplicative effects. For the AML and T2D data, we included class membership as a factor in the design matrix when applying ComBat and RBE.

#### AML components and variables

For benchmarking, we fitted models with one component because tuning indicated that performance declined with two or more components (Section 4.1.5, Supplementary Figures S4–S5). The models selected 50 accessibility variables, 50 methylation variables, and 50 RNA variables. For the final MINT Block model, we computed two components for visual comparison with the PCA plots, but focused on the first component only for interpretation, selecting 50 accessibility, 75 methylation, and 75 RNA variables for each component based on tuning (Section 4.1.6).

#### GBR components and variables

For benchmarking, we fitted models with two components based on previous MINT analyses in [36], and selected 50 taxonomic variables and 50 functional variables for each component. Tuning (Supplementary Figure S6) indicated that one component with 50 taxonomic variables, 50 functional variables, and five environmental variables was optimal. However, for the detailed MINT Block analysis, we fitted two components, each selecting 25 taxonomic variables, 25 functional variables, and five environmental variables, because this yielded a more informative biological interpretation.

#### T2D components and variables

For benchmarking, we fitted models with two components, selecting 50 species variables and 50 pathway variables for each component, as tuning indicated that performance declined with three or more components (Supplementary Figures S7–S8). Because preliminary analysis showed low predictive performance, we set the *design* argument to 0.10. Setting the *design* argument to 0.10 changed *c*_q,k_ in equations (2) and (12) from 1 to 0.10 for connections between explanatory blocks (*q, k < Q*), thereby directing MINT Block, Block (s)PLS, and DIABLO algorithms to prioritise covariances between explanatory components and response components. Following variable tuning, the final MINT Block model selected 50 species variables and 25 pathway variables for component 1, and 75 variables from each block for component 2.

### 4.6 Evaluation methods

#### ARI

The Adjusted Rand Index (ARI) [73] measures the agreement between two different partitions, with an expected value of 0 for random partitions and a maximum value of 1 indicating perfect agreement. For a given model fit and block, we applied k-means clustering [74] to the component scores, setting the number of clusters equal to the number of studies. We then calculated the ARI between the resulting clusters and study membership.

#### LOGOCV

Leave-One-Group-Out Cross-Validation involves holding out one study at a time, training the model on the remaining studies, and then predicting the held-out study. For regression, we predicted the response variables; for classification, we predicted the class of each held-out observation [22].

#### Stability

We calculated variable-selection stability from the frequency with which variables were selected during LOGOCV. We define the stability score as

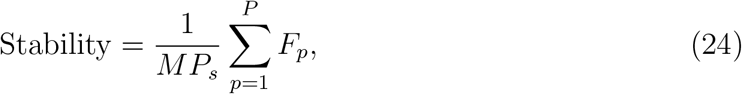

where *M* is the number of studies, *P* is the total number of variables, *P*_s_ is the number of variables selected at least once, and *F*_p_ ≤ *M* is the frequency with which the *p*th variable was selected. The stability score can range from 1*/M* (no variable selected more than once) to 1 (the same variables selected in every model fit).

#### BER

The balanced error rate (BER) for *K* classes is defined as

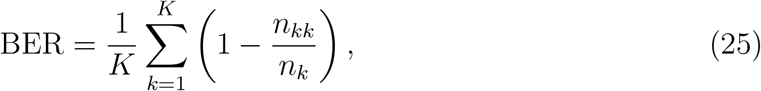

where *n*_k_ is the number of observations in class *k* and *n*_kk_ is the number of correctly classified observations in that class. Thus, each class contributes equally to the BER, regardless of its sample size [48]. For the AML and T2D models, we predicted the class of held-out observations during LOGOCV and averaged the BER across studies to calculate the overall BER.

#### RMSE

For the GBR case study, we predicted the response variables using LOGOCV. To account for differences in how the models scale input data, we centred and scaled the observed and predicted values for each variable before computing RMSE values. Let 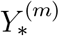 and 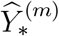 denote these centred and scaled matrices for sampling trip *m*. We calculated

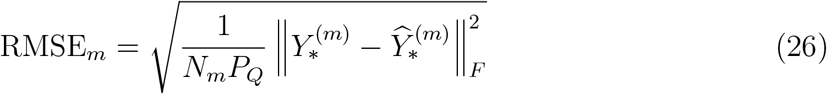

for each sampling trip (Supplementary Figure S1) and averaged these values across trips. We also computed RMSE by variable and trip, and averaged across trips to obtain RMSE values by variable (Supplementary Figure S2).

#### GO terms over-representation analysis

We used the *enricher* function from the *clusterProfiler* R package [75] to perform over-representation analysis of RNA variables selected with MINT Block, compared to a gene universe consisting of all RNA variables included in the model. Using a *p*-value cutoff of 0.01, we tested for over-representation of GO Biological Process terms [76] containing between 5 and 250 genes, obtained with the *msigdbr* R package, and plotted up to 20 of the most statistically significant terms using the *treeplot* function from the *enrichplot* R package. Terms in the *treeplot* were clustered using Jaccard similarity coefficients calculated by the *pairwise termsim* function.

#### Correlations between selected variables

To quantify and visualise associations between variables, we used pairwise similarity scores calculated with the *mixOmics network* function [77]. These similarity scores, which are the inner products of projections of variables onto components, are analogous to correlation coefficients.

## 5 Declarations

### Ethics approval and consent to participate

Not applicable.

### Consent for publication

Not applicable.

### Availability of data and materials

MINT Block is implemented in the *mixOmics* [48] R package.

### Authors’ contributions

KALC and DPH conceived the project. DPH, MT, and SMB generated and pre-processed the data. DPH conceived and implemented the analyses with input from all authors. KALC identified the case studies, developed the original version of the method and supervised the project. All authors read, edited and approved the final manuscript.

## Acknowledgments

We would like to thank Dr Yiwen Wang and Pei-Hsiu Kao for advice on analysing the T2D data.

## Supplementary Material

### S1 Supplementary text

#### S1.1 Detailed interpretation of the AML case study

To assess the biological relevance of our fitted model, we surveyed published evidence linking RNA variables selected by MINT Block to AML or cancer. For a concise overview, we focused on the top 25 variables ranked by the magnitude of their component 1 loading (Figure 2f).

Eighteen genes had higher average expression in the AZA treatment group. RBM5 [1], NAMPT [2] and NCOA7 [3] have reported roles in AML cell survival. REL encodes a subunit of NF-kappa-B which is associated with AML [4], and IL13RA1 is associated with IL4 which is thought to induce apoptosis of AML cells [5]. POLB has been associated with the prevention of tumorigenesis [6], YEATS2 has recently been identified as a potential target for cancer therapy [7]. PLK3 [8, 9] and FOCAD [10, 11] are considered tumour suppressors, while KDSR is associated with leukemia cell maintenance [12]. CAPN1 [13, 14, 15], CMTR1 [16], VOPP1 [17, 18], ATAD2B [19], and PDZD8 [20] have all been linked to multiple types of cancer.

Seven genes had higher average expression in the DAC treatment group, of which CYP1B1 has previously been linked to AML progression [21] and RHEX has been identified as a potential therapeutic target for AML [22, 23]. Among the remaining DAC genes, ZNF595 has been linked to gastric cancer [24], PNMA5 is thought to play a role in promoting carcinogenesis and progression in colorectal cancer [25], ANXA2 is linked to multiple cancer types through its influence on the migration and proliferation of cancer cells [26, 27], and MAP3K11 is believed to function as a tumour suppressor [28].

We identified GO biological process terms over-represented among all RNA variables selected for component 1 (Section 4.6, see Figure 2g), and found that many had previously been associated with AML or cancer more broadly (Table S4). Over-represented GO terms with an existing AML association included biological processes related to cell death that have been identified as potential targets for treating AML [29, 30]; processes related to responses to oxidative stress, which have been associated with AML pathogenesis and treatment [31]; respiratory system development potentially linked to respiratory complications in AML [32]; and processes involved in the regulation of Jun kinase activity [33] and T cell receptor signaling [34]. Among the remaining GO terms, those previously linked to cancer in general included biological processes related to glutamate receptor signalling [35, 36], DNA repair and organisation [37, 38], organization of intermediate filaments [39] and cellular interaction with copper ions [40].

#### S1.2 Detailed interpretation of the GBR case study

The strong trip-associated structure observed in the original data has an established ecological basis. This is consistent with well-documented seasonal and spatial variation in seawater microbiomes and water chemistry across coral reefs, including the GBR. During the wet season, greater picoplankton productivity is associated with increased particulate organic matter (POM) production, and high chlorophyll *a* concentrations, a proxy of phytoplankton biomass) [41, 42, 43]. While these patterns reflect genuine large-scale environmental and biological differences across the GBR, because each reef was sampled only once across different regions and seasons, such dominant spatio-temporal effects may mask finer-scale microbial–environment relationships. Therefore, our NP-integration with MINT Block had two objectives: first, to identify correlated multi-omics signatures by integrating taxonomic and functional omics blocks with seawater physico-chemical measurements (N-integration); and second, to assess whether these signatures were reproducible across independent spatio-temporal sampling trips (P-integration).

The opposing relationships between dissolved 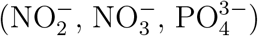 and particulate (POC, PN, PP) nutrient pools likely reflect the coupling between nutrient drawdown and biological production, consistent with ecological frameworks proposed in [44]. The microbial taxa with the strongest positive associations with particulate organic matter (POM), and negative associations with dissolved inorganic nitrogen (DIN), were dominated by picocyanobacteria from the order *Synechococcales*, including *Synechococcus, Cyanobium*, and *Parasynechococcus* (Figure 3e), while *Prochlorococcus* showed comparatively weaker associations. This is consistent with the proposed use of increased *Synechococcus*:*Prochlorococcus* abundance ratios as indicators of nutrient enrichment in coral reefs, including the GBR [43]. These associations are biologically plausible because cyanobacterial photosynthetic production contributes to phytoplankton-derived particulate organic matter in offshore GBR waters [41, 45, 46]. This interpretation is supported by the positive association between elevated chlorophyll *a* concentrations and POM (Figure 3d). Conversely, negative correlations between DIN and POM may reflect the uptake and depletion of dissolved inorganic nutrients during picocyanobacterial growth.

Newly produced organic matter would support heterotrophic microbial taxa involved in particle colonisation and remineralisation as part of the ‘microbial loop’ (proposed in [44]), including members of *Flavobacteriales, Cryomorphaceae, Rhodobacterales*, and *Marinovum*, which are known to metabolise phytoplankton-derived organic substrates [42]. The selected functional variables were also consistent with increased microbial growth and metabolic activity, including core metabolism, respiration, ATP generation and biosynthetic functions involving nucleotide synthesis, NAD(P)-dependent redox enzymes, carbamoyl-phosphate synthases and ATP synthase subunits. Together, these taxonomic and functional patterns suggest that MINT Block recovered coordinated microbial responses associated with biological productivity and organic-matter utilisation across GBR reefs (Figure 3d, e).

#### S1.3 Detailed interpretation of the T2D case study

Among the pathway variables selected for component 1, the strongest contributors to discrimination had higher average abundance in the healthy samples (Figure 4e). These included pathways involved in Colanic acid building block biosynthesis, CMP-legionaminate biosynthesis, GDP-mannose related processes, and the degradation of *β*D-glucuronide and D-glucuronate, which have previously been identified as diabetes biomarkers in humans and dogs [47, 48, 49, 50] (Table S5). Several additional pathways with higher average abundance in healthy samples were not themselves established diabetes biomarkers, but have been associated with measures related to diabetes risks of metabolic health. These included pathways involved in the degradation of D-galacturonate, hexuronide, hexuronate, 4-deoxy-L-threo-hex-4-enopyranuronate, and D-fructuronate, with reported associations involving gamma-glutamyl transpeptidase levels [51, 52], non-high-density lipoprotein cholesterol levels [53, 54], production of short fatty acids [55, 56], and the dietary inflammatory pattern score [57]. This group also included a glutaryl-CoA degratation pathway associated with metabolic disorders[58], and *β*-(1-4)-mannan degradation, which is altered by dietary interventions in pre-diabetic patients [59].

Among the pathways with higher average abundance in T2D samples (Table S6), several were involved in purine metabolism, which has been associated with diabetes progression [60]. Related pathways have also been reported in diabetic dogs [49] and children of mothers with gestational diabetes [61].

Other pathways with higher average abundance in T2D samples were involved in methanogenesis and pyrimidine deoxyribonucleotides biosynthesis that have been linked to obesity [62], and pathways involved in L-lysine biosynthesis. Although L-lysine biosynthesis is not an established diabetes biomarker, altered lysine metabolism has been reported in diabetic rats [63]. The superpathway of L-serine and glycine biosynthesis I also had a higher average abundance in the T2D group. Because lower glycine and L-serine metabolite levels have previously been reported in T2D, the biological interpretation of this pathway association remains uncertain and may reflect either a compensatory microbial response or a spurious association [64, 65].

We next examined the correlations between the top 25 species and pathway variables selected for component 1, ranked by absolute loading. The selected species formed three subsets based on their abundance patterns and pathway correlations (Figure 4f, see Section 4.6 for how correlations were calculated). The first subset comprised species with higher average abundance in healthy samples: *Lachnospira eligens, Ruminococcus torques, Ruminococcus bicirculans, Intestinibacter bartlettii, Proteobacteria bacterium CAG:139, Turicimonas muris, Haemophilus parainfluenzae, Romboutsia ilealis, Ruminococcus lactaris*, and *Eubacterium sp. CAG:38*. These species were positively correlated with most of the pathways with higher average abundance in healthy samples and reported links to diabetes or metabolic health, particularly pathways involving cell-surface polysaccharide biosynthesis, uronic-acid degradation, GDP-mannose biosynthesis and glutaryl-CoA metabolism (Table S5).

The second subset comprised *Enterocloster lavalensis, Blautia sp. CAG:257*, and *Clostridium symbiosum*, which had higher abundance in the T2D group, and *Enterocloster clostridioformis* which was higher in the healthy group despite having a component loading in the T2D direction. These species were positively correlated with T2D-associated pathways involved in purine metabolism (purine nucleobases degradation), L-serine and glycine biosynthesis, and L-lysine biosynthesis, that are associated with diabetes or metabolic health (Table S6).

The third subset, which also had higher abundance in the T2D group included *Tractidigestivibacter scatoligenes, Collinsella aerofaciens, Slackia isoflavoniconvertens, Collinsella intestinalis, Enorma massiliensis, Clostridium leptum*, and *Streptococcus vestibularis*. These species were positively correlated with T2D-associated pathways involving pyrimidine deoxyribonucleotides biosynthesis, L-lysine biosynthesis, methanogenesis, and purine metabolism (Inosine-5’-phosphate biosynthesis) (Table S6).

### S2 Supplementary tables

**Table S1:** AML cell counts by treatment group and batch.

| Treatment | Batch |  |  |
| --- | --- | --- | --- |
|  | b320 | b322 | b620 |
| AZA | 20 | 34 | 37 |
| DAC | 21 | 36 | 36 |

**Table S2:** GBR sample counts by sampling trip and GBR sector.

| Trip | GBR sector |  |  |
| --- | --- | --- | --- |
|  | North | Central | South |
| Trip 01 Nov-Dec 2019 | 44 | 0 | 0 |
| Trip 02 Jan 2020 | 0 | 0 | 48 |
| Trip 03 Feb 2020 | 0 | 43 | 0 |
| Trip 04 Jul 2020 | 0 | 56 | 0 |

**Table S3:**
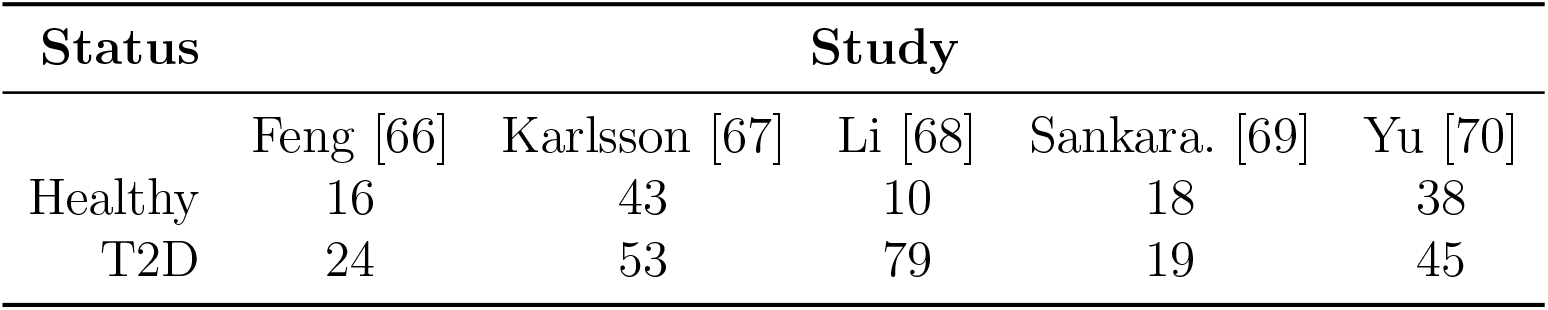
T2D sample counts by status and study.

**Table S4:**
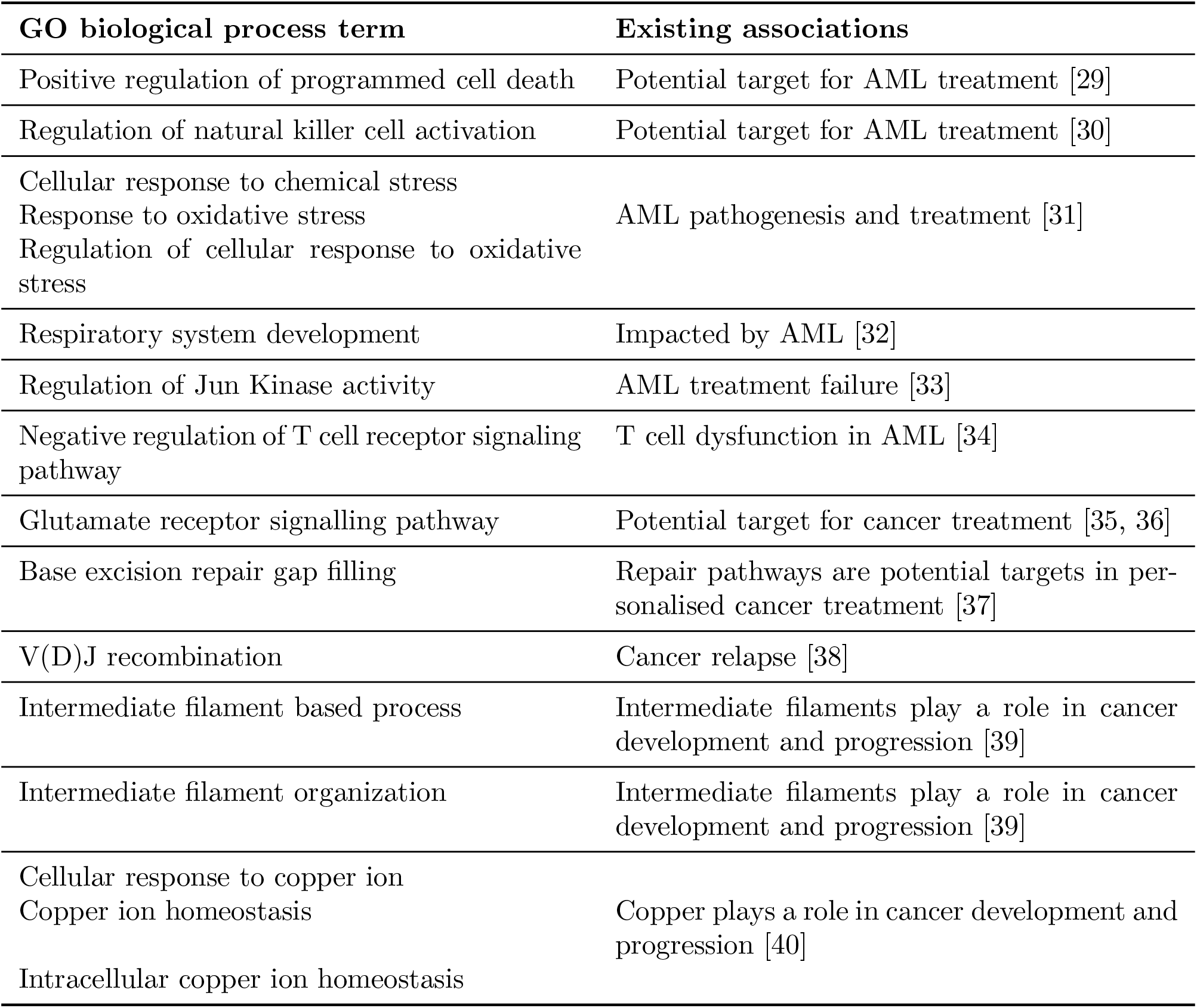
GO biological processes over-represented among RNA variables selected on MINT Block component 1 in AML cells. The enriched terms include processes previously associated with AML pathogenesis or treatment and broader cancer-related mechanisms involving DNA repair, intermediate-filament organisation and copper homeostasis.

**Table S5:** T2D case study. Metabolic pathways with higher average abundance in healthy samples and selected on MINT Block component 1. Several have reported associations with diabetes risk or metabolic health, including pathways involving carbohydrate degradation, GDP-mannose and colanic acid biosynthesis, and glutaryl-CoA metabolism.

| Pathway ID | Name | Existing associations |
| --- | --- | --- |
| GALACTUROCAT-PWY | D-galacturonate degradation I | Associated with insulin levels [71] (metabolic health) and gamma-glutamyl transpeptidase (GGT) levels [51] (potential biomarker for diabetes risk [52]) |
| GALACT-GLUCUROCAT-PWY | Superpathway of hexuronide and hexuronate degradation | Associated with Non-high-density lipoprotein cholesterol levels [53] (potential biomarker for diabetes risk [54]) and GGT levels [51] (potential biomarker for diabetes risk [52]) |
| PWY-6507 | 4-deoxy-L-threo-hex-4-enopyranuronate degradation | Involved in the production of short fatty acids [55], which potentially protect against diabetes [56]; associated with GGT levels [51] (potential biomarker for diabetes risk [52]) |
| PWY-5659 | GDP-mannose biosynthesis | Diabetes biomarker in humans [47] and dogs [49]; associated with GGT levels [51] (potential biomarker for diabetes risk [52]) |
| GLUCUROCAT-PWY | Superpathway of βD-glucuronide and D-glucuronate degradation | Depleted in diabetes patients [50]; associated with GGT levels [51] (potential biomarker for diabetes risk [52]) |
| COLANSYN-PWY | Colanic acid building blocks biosynthesis | Diabetes biomarker [47]; linked to protection against gestational diabetes mellitus [72]; associated with GGT levels [51] (potential biomarker for diabetes risk [52]) |
| PWY-7242 | D-fructuronate degradation | Associated with the empirical dietary inflammatory pattern score [57] (diabetes risk) |
| PWY-7323 | Superpathway of GDP-mannose-derived O-antigen building blocks biosynthesis | Diabetes biomarker in dogs [49] |
| PWY-5177 | Glutaryl-CoA degradation | Involved in glutarylation which is linked to the metabolic disorder glutaracidemia type I [58] |
| PWY-7456 | β-(1-4)-mannan degradation | Impacted by dietary interventions in pre-diabetic patients [59] |
| PWY-6749 | CMP-legionaminate biosynthesis I | Diabetes biomarker [48] |

**Table S6:** T2D case study. Metabolic pathways with higher average abundance in T2D samples and selected on MINT Block component 1. These include pathways involved in purine and pyrimidine metabolism, methanogenesis and amino-acid biosynthesis, which have reported associations with diabetes or related metabolic traits.

| Pathway ID | Name | Existing associations |
| --- | --- | --- |
| P164-PWY | Purine nucleobases degradation I (anaerobic) | Involved in purine metabolism which is associated with diabetes [60]; biomarker for diabetes in dogs [49] |
| PWY-6123 | Inosine-5'-phosphate biosynthesis I | Involved in purine metabolism which is associated with diabetes [60]; enriched in children of mothers with gestational diabetes mellitus [61] |
| PWY-6124 | Inosine-5'-phosphate biosynthesis II | Involved in purine metabolism which is associated with diabetes [60] |
| PWY-7234 | Inosine-5'-phosphate biosynthesis III | Involved in purine metabolism which is associated with diabetes [60] |
| METH-ACETATE-PWY | methanogenesis from acetate | Involved in methane production (glucose tolerance and susceptibility to hyperglycemia [73]) and GLP-1 secretion [74]; associated with obesity in children [62] |
| PWY-7187 | Pyrimidine deoxyribonucleotides de novo biosynthesis II | Associated with obesity in children [62] |
| PWY-7198 | Pyrimidine deoxyribonucleotides de novo biosynthesis IV | Associated with obesity in children [62] |
| PWY-2941 | L-lysine biosynthesis II | Lysine degradation enriched in the kidneys of diabetic rats [63] |
| PWY-2942 | L-lysine biosynthesis III | Lysine degradation enriched in the kidneys of diabetic rats [63] |
| SER-GLYSYN-PWY | Superpathway of L-serine and glycine biosynthesis I | L-serine [65] and Glycine [64] levels depleted in T2D |

### S3 Supplementary figures

**Figure S1:**
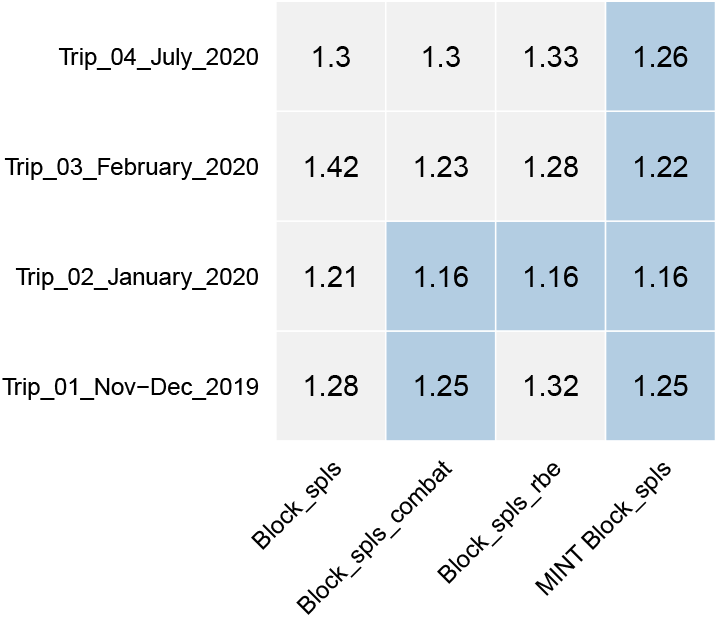
GBR RMSE values by sampling trip for the different models evaluated during benchmarking. The lowest RMSE for each trip is highlighted in blue. For each trip and fitted model, the RMSE was calculated as the square root of the mean squared prediction error across all elements of the response matrix. MINT Block achieved the lowest RMSE, including ties, for every trip.

**Figure S2:**
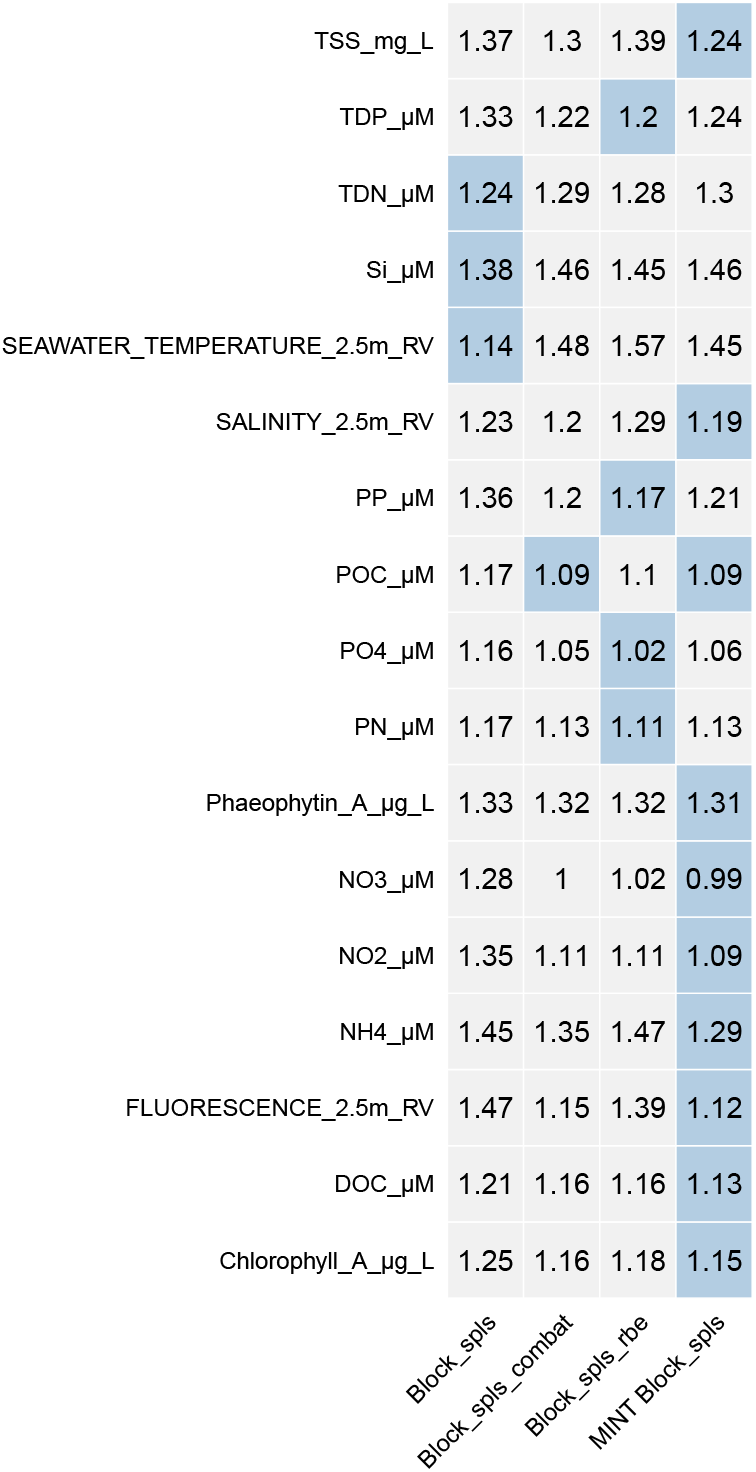
GBR RMSE values by response variable for the different models evaluated during bench-marking. The lowest RMSE for each variable is highlighted in blue. The RMSE was first calculated for each response variable, fitted model and trip, and then averaged across the trips. MINT Block achieved the lowest RMSE, including ties, for 10 out of 17 response variables.

**Figure S3:**
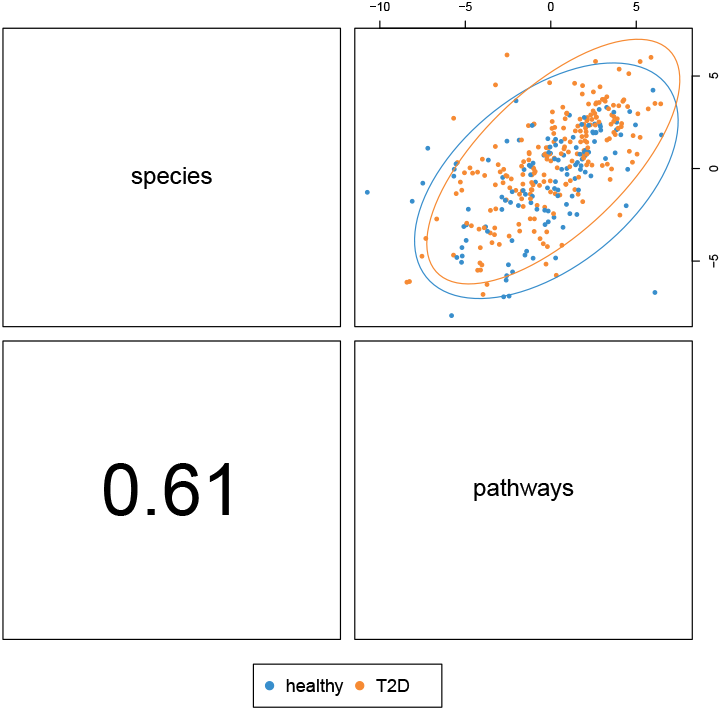
T2D sample plot for MINT Block component 2, coloured by T2D status. Component 2 showed little discrimination by T2D status but stonger correlations between blocks than component 1.

**Figure S4:**
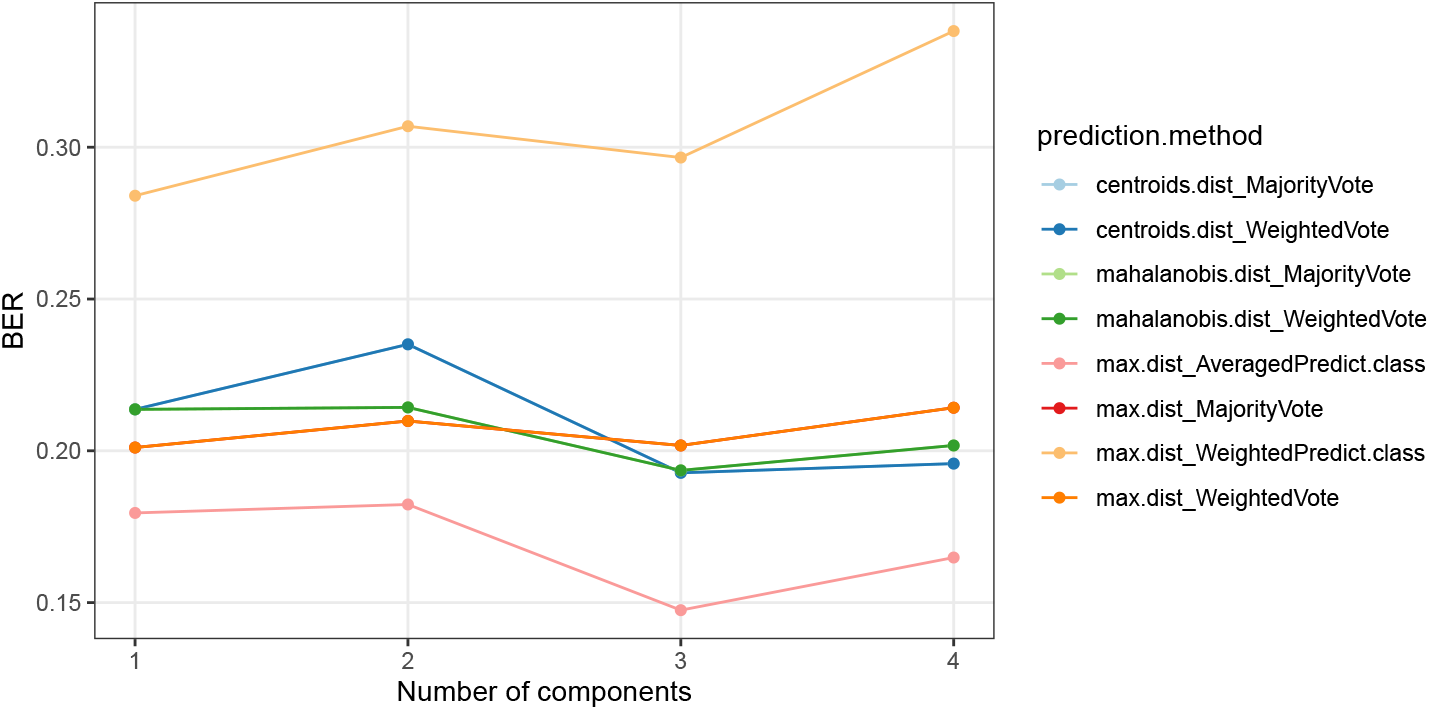
AML BER values for MINT Block obtained using LOGOCV, plotted by prediction method and number of components. The lowest BER was achieved by the classification method that averages response values predicted from each explanatory block.

**Figure S5:**
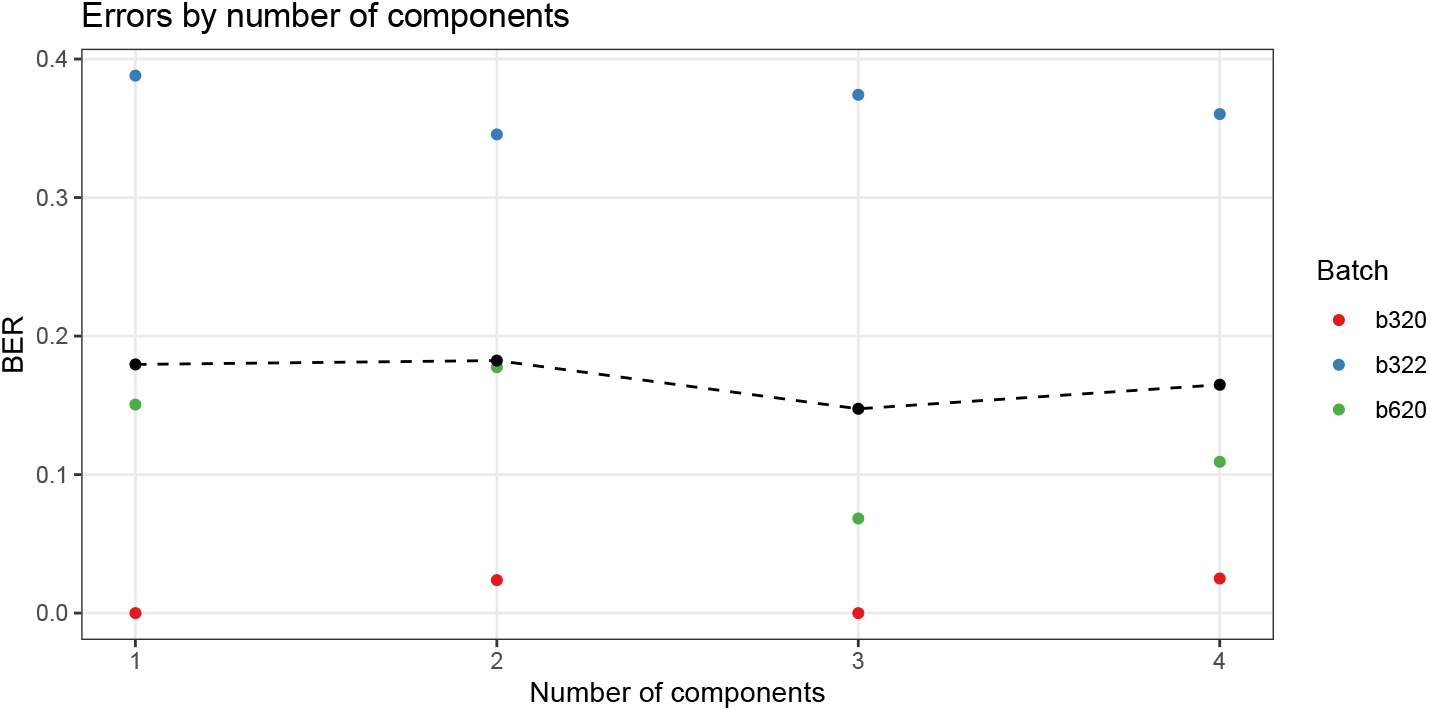
AML BER values for MINT Block using LOGOCV, plotted by study and number of components using the prediction method with the lowest overall BER. The black dots represent the mean BER across studies for each number of components. Using more than one component did not improve prediction accuracy.

**Figure S6:**
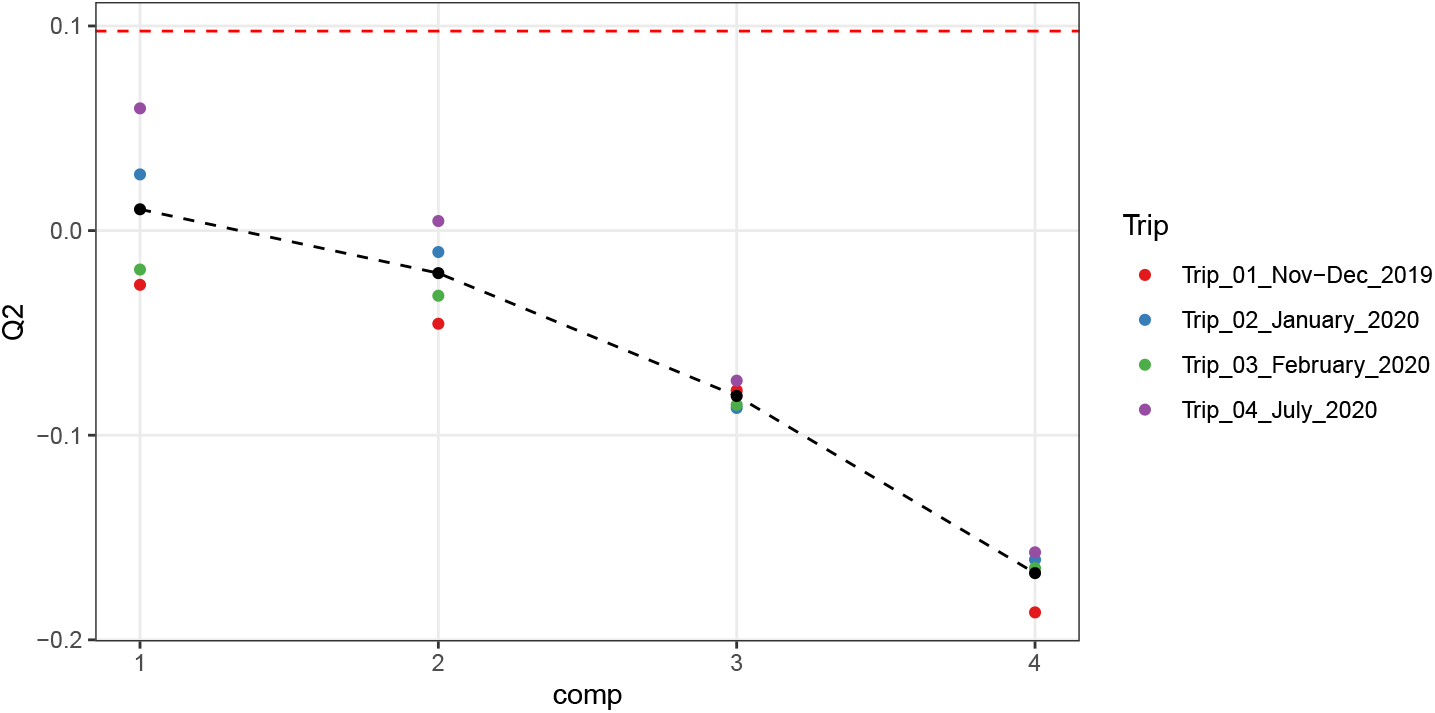
GBR *Q*^2^ values for for MINT Block using LOGOCV, plotted by study and number of components. The black dots represent the mean for each number of components. Using more than one component did not improve prediction accuracy.

**Figure S7:**
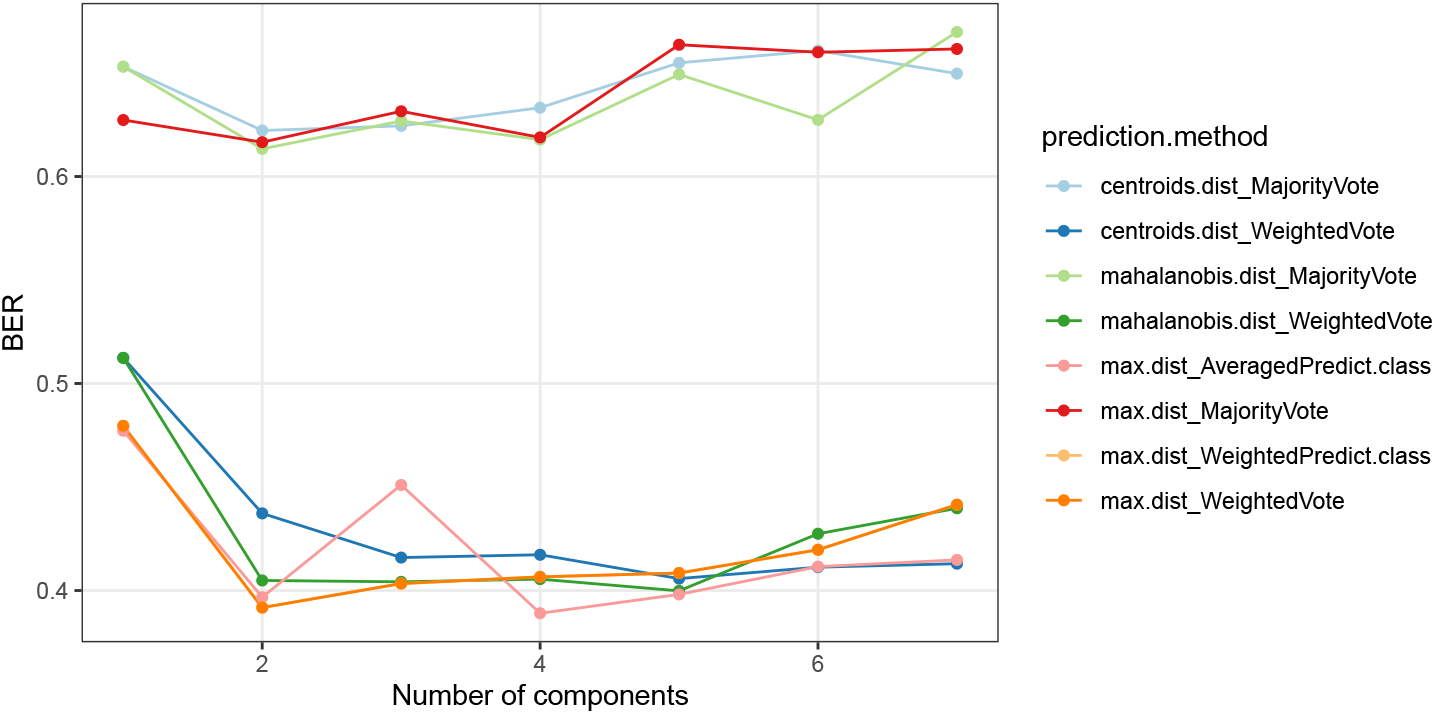
T2D BER values for MINT Block using LOGOCV, plotted by prediction method and number of components. Most prediction methods, except those based on majority voting across the explanatory blocks, performed similarly.

**Figure S8:**
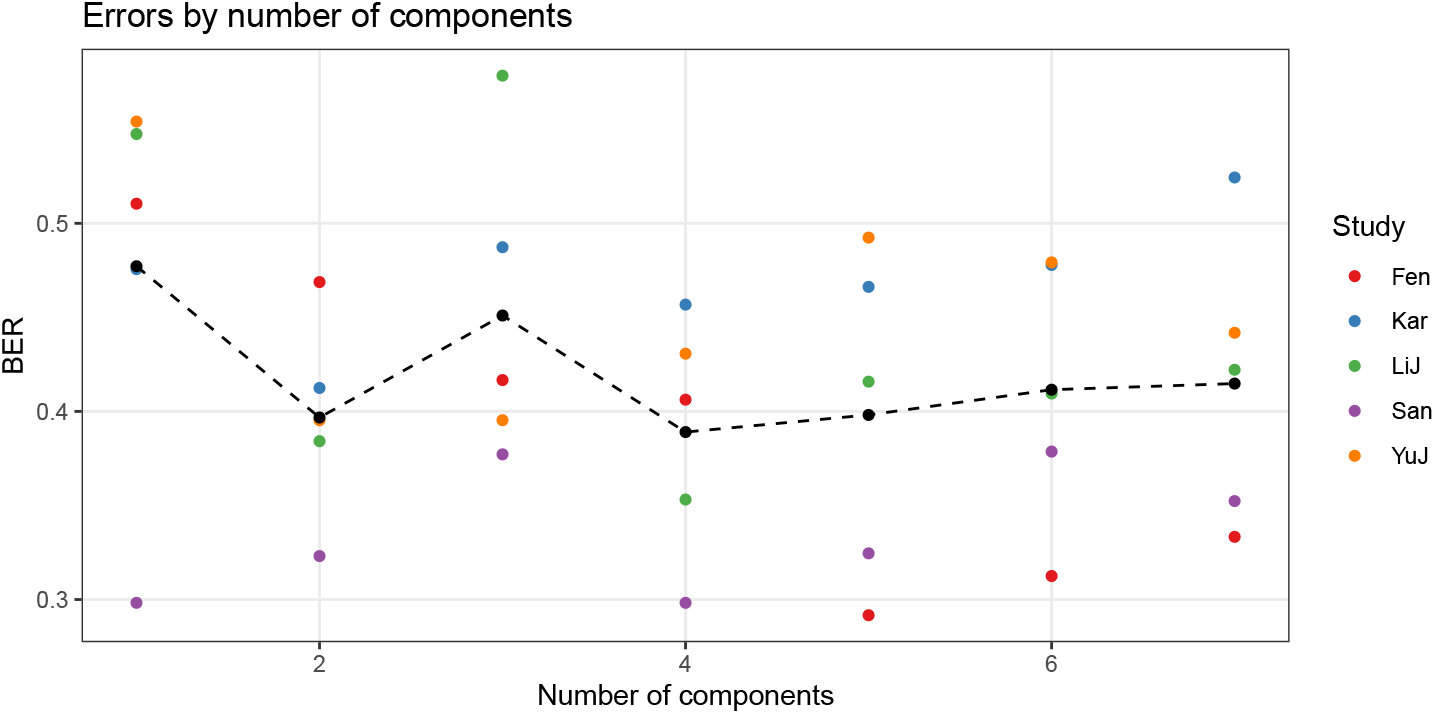
T2D BER values for MINT Block using LOGOCV, plotted by study and number of components using the prediction method with the lowest overall BER. The black dots represent the mean for each number of components. Prediction accuracy improved when the number of components was increased from one to two, and then plateaued.

